# ZHP-2/HEI10 generates CO precursors and triggers procrossover factor coarsening and CO formation through Polo kinase recruitment

**DOI:** 10.64898/2026.07.26.740636

**Authors:** Alyssa Centonza-Brenie, Ryan Dawson, Sowmya Geetha, Aya Sato-Carlton, Peter M. Carlton, Verena Jantsch, Monique Zetka

**Affiliations:** Department of Biology, McGill University, Montreal, H3A 1B1 Canada; Max Perutz Laboratories, Vienna Biocenter Campus, 1030, Vienna, Austria; University of Vienna, Vienna Biocenter Campus, 1030, Vienna, Austria; Graduate School of Biostudies, Kyoto University, Yoshida-Konoecho, Kyoto 606-8501, Japan; Radiation Biology Center, Kyoto University, Yoshida-Konoecho, Kyoto, 606-8501, Japan

**Author notes:** Equal contribution (Alyssa Centonza-Brenie, Ryan Dawson).

## Abstract

During meiosis, programmed DNA double-strand breaks (DSBs) are repaired with the homolog to form crossovers (COs) essential for chromosome segregation, or noncrossovers (nCOs) to restore genomic integrity. Only a few recombination intermediates will become COs and these sites are progressively enriched in proCO factors at the expense of nCO sites, referred to as coarsening. Here, we show that in *C. elegans* this process is engineered by ZHP-2/HEI10, a conserved E3 ligase that has functions at early and late stages of CO formation. Early on, ZHP-2-mediated loss of RMH-1/RMI1 and HIM-6/BLM from strand exchange intermediates is required for joint molecule formation and nCO repair. In *zhp-2* mutants, proCO factors accumulate at HR sites, leading to defects in CO formation and CO spacing. We find that Polo kinase recruitment to a ZHP-2 at CO precursors signals an end to further DSB formation, and triggers proCO factor coarsening and CO formation. We propose that ZHP-2 disassembles proCO factors from recombination intermediates until it is inactivated at CO precursors through Polo kinase recruitment, leading to stabilization of proCO factors, and resolution of joint molecules as crossovers. Our study reveals that ZHP-2/HEI10 is the central component of a Polo kinase-driven switch that initiates proCO factor coarsening and execution of crossing over.

## Introduction

During meiosis, DNA double-strand breaks (DSBs) are induced as a means of initiating homologous recombination (HR) and physically connecting homologs through crossovers (COs). Meiotic DSBs are induced in greater numbers than the final number of COs, thereby ensuring that at least one DSB per homolog is repaired as a CO while the remainder are repaired as noncrossovers (nCOs) to restore genomic integrity. DSB formation during meiosis is catalyzed by the topoisomerase-like Spo11 protein,^1^ yielding two ends that are resected to form 3’ overhangs that invade the sister chromatid or the homolog in search of homologous sequences to use as repair templates. This process of template invasion creates D-loops structures, recombination intermediates that primarily undergo disassembly and are repaired as nCOs using synthesis-dependent strand annealing (SDSA). Alternatively, a limited number of D-loops form stable single-end invasion intermediates (SEI) where the “capture “of the second end of the resected DSB results in joint molecules (JM) that can be processed to yield double-Holliday junctions (dHJs).^2,3^ dHJs are the key precursors of COs, and their stabilization favours resolution by structure-specific endonucleases into CO products.^4,5^ As a result, the repair of meiotic DSBs has critical junctures during homologous recombination that lead a crossover outcome: second end capture to generate dHJs, and stabilization of dHJs to generate COs.

HR-mediated DSB repair can readily occur between sister chromatids,^6–9^ and to ensure CO formation, HR is biased towards using the chromatids of the homolog as a repair template.^2,10–16^ A key mediator of the interhomolog bias during DSB repair is the synaptonemal complex (SC), a conserved protein network that forms between homologous chromosome axes and is required for CO formation.^17^ Recombination between sister chromatids is also actively discouraged by conserved helicases that disrupt D-loop strand exchange intermediates during the early steps of HR. In mammals, the Bloom syndrome helicase BLM interacts with topoisomerase IIIα (TOP3A) and the scaffolds RMI1 and RMI2 to form the BTR complex; BTR has well-documented mitotic activities in dissolving D-loop structures and sister chromatid crossover intermediates.^18,19^ During yeast meiosis, Sgs1/BLM complexes with Top3 and Rmi1 to similarly regulate HR through dissolution of D-loop structures; this has the effect of preventing unwanted HR intermediates between chromatids, and recycling of the DSB ends.^6,20–22^ The latter has led to the proposal that when one end is stabilized by Msh4-Msh5, the other end is eventually freed by Sgs1/BLM for another attempt at second end capture and joint molecule formation.^6^ In *C. elegans*, HIM-6/BLM and RMH-1/RMI1 first appear at early strand exchange HR intermediates and then concentrate at late prophase CO sites, evidence of a pro-CO activity that is distinct from an earlier function in D-loop disruption.^23,24^

One of the least understood aspects of meiosis is how one recombination intermediate among many is assigned the CO fate. A conserved cytological feature of meiosis of diverse organisms is the emergence of a few prominent CO structures from a much larger pool of smaller recombination intermediates that appear earlier.^25,26^ Recombination intermediate dynamics are visible in the *C. elegans* germline, where a procession of conserved proCO factors initially appear as numerous small foci that dissipate from future nCOs sites and accumulate at the few sites designated to become COs: MSH-5/MutSγ, the checkpoint kinase CDK-2 with its cyclin COSA-1/CNTD1, and the HIM-6/BLM helicase with its scaffold RMH-1/RMI1.^23,27–38^ The progressive concentration of proCO factors at the sites destined to become COs is critical for stabilizing late CO intermediates like dHJs, and fostering their resolution into COs at late prophase.^24,33,38,39^

In diverse organisms, the restriction of proCO factors to CO sites is regulated by a family of conserved SUMO/Ubiquitin E3 ligases that function in the CO/nCO decision.^40^ In mice for example, the SUMO E3 ligase RNF212 localizes throughout the SC and is then confined to a few foci where it stabilizes MutSγ, leading to CO formation.^41^ The restriction of RNF212 to CO sites depends on HEI10, a ubiquitin ligase that similarly first localizes to the SC and then to the sites of crossing over;^42^ in the absence of HEI10, RNF212 (and MutSγ) are not restricted, and COs do not form.^43^ The analysis of meiotic RING finger proteins in Sordaria, plants, mice, and *C. elegans* has led to the proposal that COs are the outcome of CO factor stabilization at future CO sites and their concomitant loss from future nCO sites.^20,41,42,44–47^ The combined results of these studies are consistent with a model in which mobile proCO factors become stabilized at a CO intermediate, leading to a positive feedback loop that results in proCO factor coarsening and CO formation.^35,40,47,48^ The progressive stabilization of proCO factors at designated CO sites is likely to represent expanding networks of protein-protein interactions that are fueled by CO factor entrapment, resulting in protection of joint molecules and their resolution into COs.

The RNF212 – HEI10 circuit required for CO formation in mammals is represented in *C. elegans* by four orthologs that interact to form two distinct SC-localized complexes (ZHP-1/2, and ZHP-3/4) that are essential for crossing over.^28,32,34^ During meiotic recombination, the ZHP complexes have antagonistic roles that drive differentiation between CO and nCO sites, similar to the competing activities of RNF212 and HEI10 during mammalian CO formation. ZHP-3/4 are required for the stabilization of proCO factors (RMH-1/HIM-6, MSH-5, COSA-1) at HR sites,^32,34^ an activity most closely resembling RNF212. In contrast, ZHP-1/2 mimics the activity of HEI10 in proCO factor removal, leading to restriction of ZHP-3/4 and other proCO factors to future CO sites (this study).^34^ Although the ZHP complexes and their orthologs are known to function in CO formation by regulating proCO factor dynamics, the underlying regulatory network linking their coarsening to CO formation is still unclear. In this study, we perform an extensive characterization of ZHP-1/2 and demonstrate that the complex has early and late functions in the HR pathway that are essential for interhomolog DSB repair, proCO factor coarsening, and CO formation. Importantly, we provide evidence that ZHP-2-mediated recruitment of PLK-2 to CO intermediates is required for coarsening of proCO factors, and the maturation of CO-designated sites into crossovers.

## Results

### CO formation requires association of the ZHP-1/2 complex with the SC

An EMS-based screen for recessive meiotic mutants^32^ recovered *zhp-2(vv84)*, a predicted null allele that introduces a nonsense mutation at the third amino acid (a.a.) of the predicted protein sequence. We subsequently used CRISPR-cas9 mutagenesis^49^ to generate a frameshift mutation in its interacting partner *zhp-1* that introduced a premature stop codon at a.a. 7 in ZHP-1 (Fig. 1A). Both *zhp-1(vv111)* and *zhp-2(vv84)* mutants individually exhibited the high levels of embryonic lethality and male offspring among surviving progeny (Fig. 1B, C) that are diagnostic of aneuploidy originating from meiotic chromosome missegregation. *C. elegans* chromosome normally undergo a single exchange, and diakinesis nuclei invariably contain 6 condensed bivalents that represent the 6 homologous chromosomes pairs joined by a single chiasma.^50–52^ In *zhp-1(vv111*) and *zhp-2(vv84*) mutants, diakinesis nuclei contained an average of 11.3 DAPI-stained structures for both mutants (*P*<0.0001 in comparison to wild type; Fig. 1D, E), indicating a severe defect in CO formation. Since the function of ZHP-1 and ZHP-2 is nonredundant and dependent on the formation of a heterodimer (referred to as ZHP-1/2)^34^, we used the *zhp-2(vv84*) mutant as a proxy to study the loss-of-function of the ZHP-1/2 complex in most of our analyses. In agreement with a previous report,^34^ these revealed that the complex is not required for chromosome pairing, synapsis, meiotic DSB formation and DSB repair, but is essential for crossing over (Fig. S1, S2). In wild type, the ZHP-1/2 heterodimer colocalizes with the SC throughout pachytene, and is dependent on the SC for its localization to chromosomes, indicating that its function in CO formation requires the SC (Fig. S3).^34^

**Figure 1.**
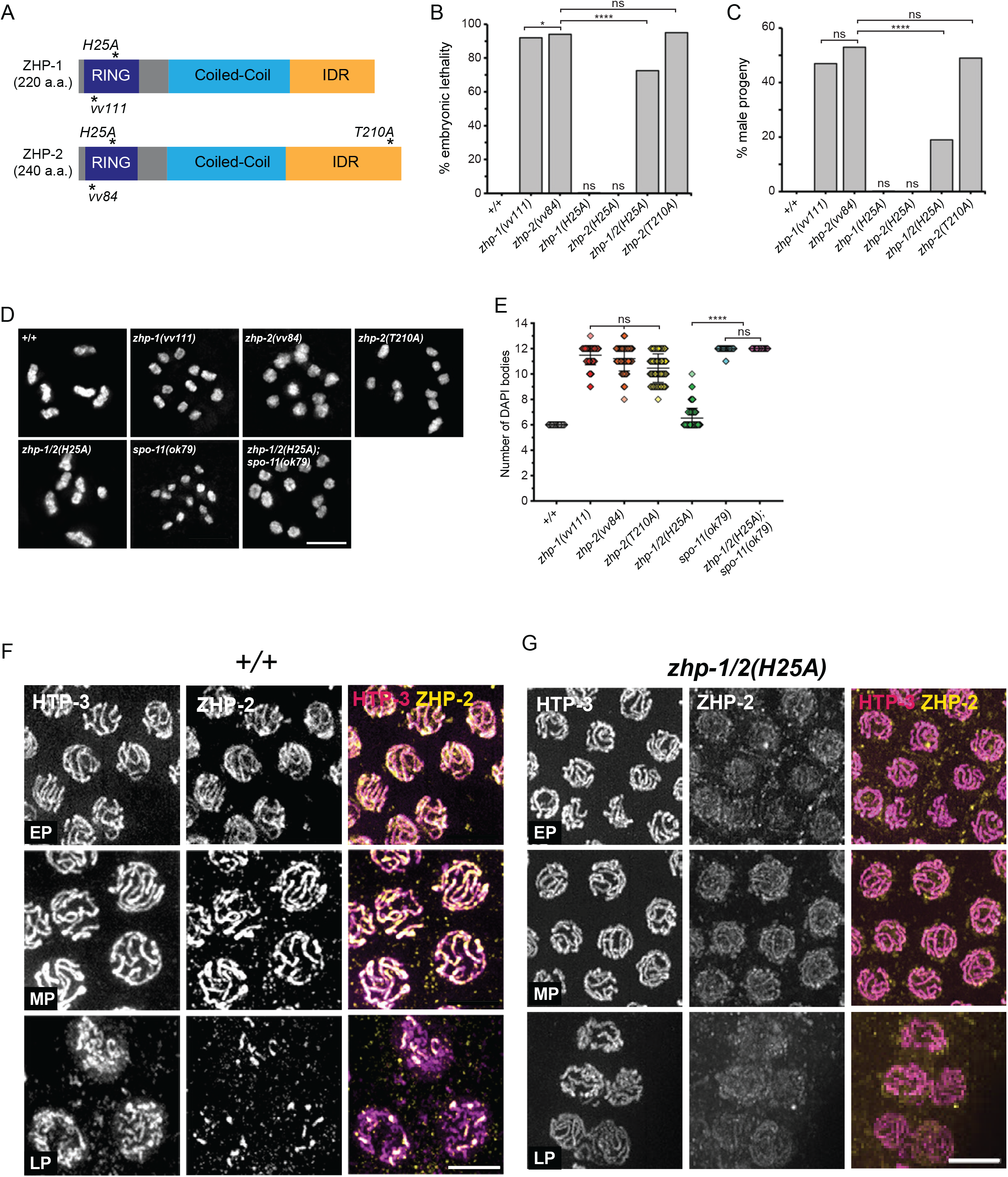
ZHP-1/2 resemble RING E3 ligases and are required for crossing over and bivalent formation. (A) Schematic of the InterPro (https://www.ebi.ac.uk/interpro/) predicted protein structures of ZHP-1 and ZHP-2. Royal blue marks the RING domain, light blue marks the coiled-coil, and orange marks the intrinsically disordered region (IDR). Locations of mutations used in this study are marked by asterisks. (B, C) Histogram graphs showing frequencies of embryonic lethality and male offspring among the progeny of the indicated genotypes. Total number of eggs scored (*n*), embryonic lethality(%)/incidence of males(%): +/+ *n*=1937, 0.2/0.11; *zhp-1(vv111*) *n*=1895, 92/47; *zhp-2(vv84) n*=1183, 94/53; *zhp-1(H25A*) *n*=1477, 0.4/0.21; *zhp-2(H25A) n*=1564, 0.13/0.13; *zhp-1/2(H25A) n*=1213, 72.5/19; *zhp-2(T210A) n*=1701, 95/49. Statistical significance was assessed by Chi-squared test followed by Bonferroni correction. Analyses revealed a statistical difference between *zhp-1/2(H25A)* mutants and *zhp-1(vv111)* or *zhp-2(vv84)* mutants (*p*<0.0001) for both embryonic lethality and male progeny. Comparisons that did not show a statistical difference for embryonic lethality/incidence of males: *+/+* to *zhp-1(H25A) P*= 0.2847/0.4483, to *zhp-2(H25A) P*= 0.5761/0.8309; *zhp-2(vv84)* to *zhp-1(vv111) P*=0.0355/0.3432, to *zhp-2(T210A) P*=0.0833/0.055. (D) Representative diakinesis nuclei from the indicated genotypes stained with DAPI. Scale bar, 5 µm. (E) Scatterplot graph showing the number of DAPI bodies in the −1 diakinesis oocytes of the indicated genotypes. Error bars denote the mean ± standard deviation. Nuclei scored (*n*), mean ± standard deviation: +/+ *n*=80, 6.0 ± 0.0; *zhp-1(vv111) n*=79, 11.5 ± 0.8; *zhp-2(vv84) n*=58, 11.2 ± 1.0; *zhp-2(T210A) n*=100, 10.46 ± 1.1; *zhp-1/2(H25A) n*=164, 6.5 ± 0.8; *spo-11(ok79) n*=74, 12.0 ± 0.2; *zhp-1/2(H25A); spo-11(ok79) n*=55, 12.0 ± 0.0. Statistical significance assessed by Kruskal-Wallis test and post Dunn’s test: ns=not significant (*P*≥0.05), **** *P*<0.0001. *zhp-1(vv111)* to *zhp-2(vv84) P*= 1, to *zhp-2(T210A) P*= 0.0143; *zhp-2(vv84)* to *zhp-2(T210A*) *P*= 0.7571; *spo-11(ok79*) to *zhp-1/2(H25A); spo-11(ok79) P*= 1. (F,G) Immunofluorescence micrographs of representative nuclei in early (EP), mid (MP), and late pachytene (LP) marked with HTP-3 (magenta) and ZHP-2 (yellow) of the indicated genotypes. Scale bars, 5 µm.

*C. elegans* ZHP proteins resemble ubiquitin E3 ligases and have a tripartite organization that includes an N-terminus RING finger domain predicted to interact with a ubiquitin E2 conjugation enzyme (Fig. 1A). Like the ZHP-3/4 complexes, ZHP-1/2 are predicted to assemble into a rod-like tetramer that spans the SC, and consists of coiled dimers that cluster the four RING domains in the centre,^47^ a configuration that is required for the *in vitro* ubiquitin ligase activity of human HEI10.^53^ To investigate the contribution of the RING domain to ZHP-1/2 function, we targeted the histidine residue known to be essential for RING finger function in ZHP-3/4 and in E3 RING proteins found in other organisms.^32,54^ In *zhp-1(H25A)* and *zhp-2(H25A)* single mutants, ZHP-2 localized with wild-type dynamics (Fig. S4) and the mutants showed no significant increase in embryonic lethality or frequency of male progeny (Fig. 1B, C), indicating that the RING domains are functionally redundant and that two intact zinc fingers are sufficient to stabilize the predicted tetramer. However, the disruption of both RING finger domains in *zhp-1(H25A) zhp-2(H25A)* double mutants (henceforth referred to as *zhp-1/2(H25A*)) resulted in a severely disordered pattern of localization, despite the presence of fully assembled SC (Fig. 1F, G, S2B). While the mutant ZHP-1/2^H25A^ complex appeared with appropriate timing at early pachytene, it formed a faint nuclear haze with fine threads that colocalized with the SC; these increased in intensity as pachytene progressed, until dissolving into a diffuse signal in late pachytene nuclei. These data reveal that while the RING domains are not essential for localization of ZHP-1/2 *per se*, they are required to establish or maintain wild-type levels of the complex at the SC.

Despite the highly disrupted localization of ZHP-1/2^H25A^, the levels of embryonic lethality and male progeny were significantly less severe than those observed in either the *zhp-1(vv111*) or *zhp-2(vv84)* null mutants (*P*<0.0001 in comparison to either null allele; Fig. 1B, C). This mitigated aneuploidy correlated with the presence of abundant bivalents in diakinesis-stage oocytes; 90% of the *zhp-1/2(H25A)* mutant nuclei contained an average of 6.5 DAPI-stained structures (Fig. 1D, E). These represented *bona fide* bivalents since loss of SPO-11 resulted in diakinesis nuclei that invariably contained 12 univalents (*P*<0.0001 in comparison to *zhp-1/2(H25A);* Fig. 1D, E). Furthermore, CO formation in *zhp-1/2(H25A)* mutants was abrogated in the absence of the proCO factors COSA-1, ZHP-4, and RMH-1, indicating that crossing over in the double RING mutant is a product of the canonical CO pathway (Fig. 2A, B). The robust CO formation that occurs in the presence of reduced levels of ZHP-1/2^H25A^ at the SC can most simply be explained by residual interactions between the disrupted RING domains or nearby regions. In this case, tetramer formation may be disfavoured and/or unstable, resulting in reduced levels that can support CO formation in regions in which it is sufficiently assembled. Although *zhp-1/2(H25A)* mutants are highly competent for crossing over, the COs that form are defective in supporting CO-triggered processes, discussed in more detail in the following sections.

**Figure 2.**
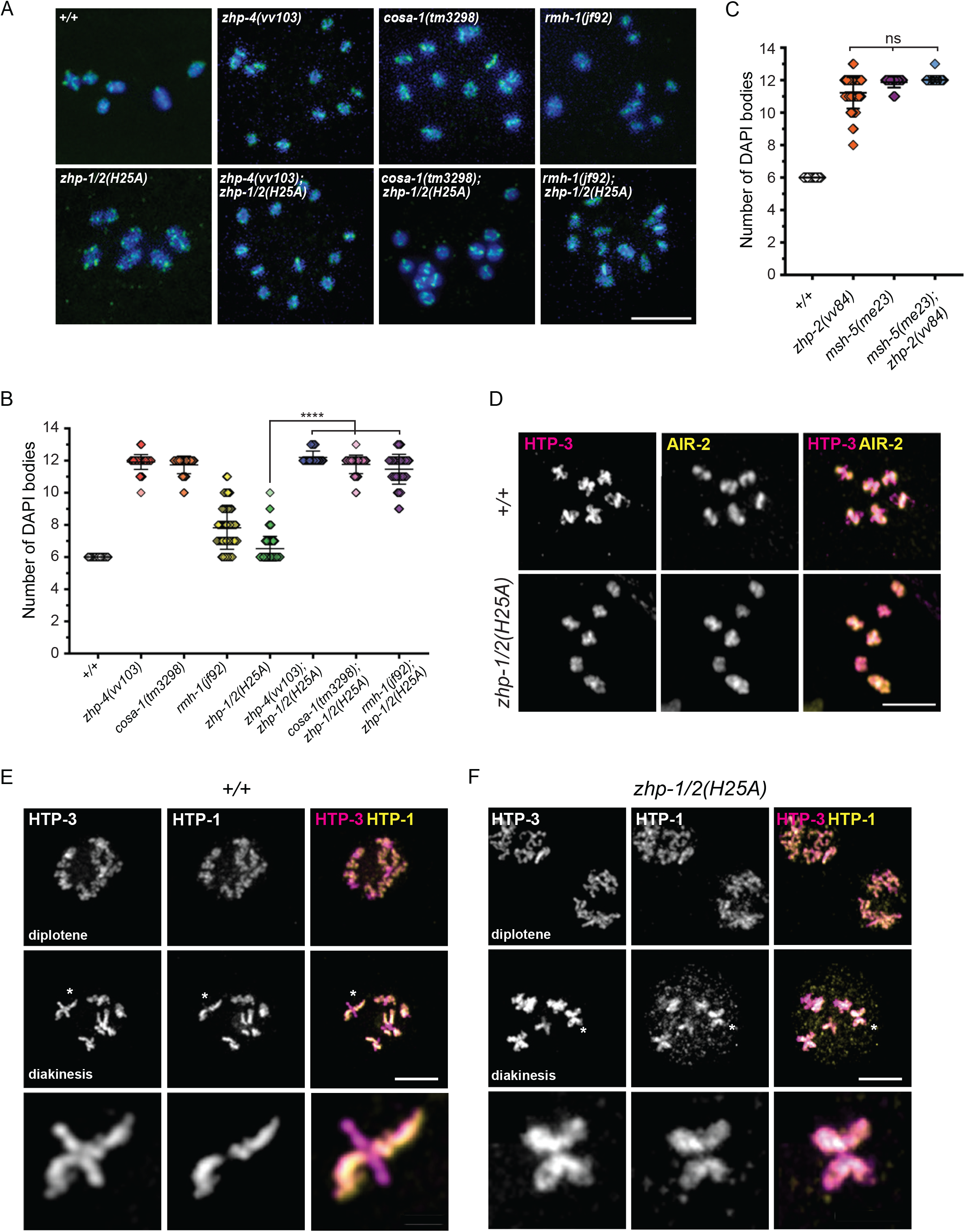
CO formation in *zhp-1/2(H25A)* mutants requires RMH-1, COSA-1 and ZHP-4. (A) Representative diakinesis nuclei from the indicated genotypes marked with DAPI (blue) and HTP-3 (green). Scale bar, 5 µm. (B) Scatterplot graph showing the number of DAPI bodies in the −1 diakinesis oocytes of the indicated genotypes. Error bars denote the mean ± standard deviation. Nuclei scored (*n*), mean ± standard deviation: +/+ *n*=30, 6.0 ± 0.0; *zhp-4(vv103) n*=78, 11.9 ± 0.5; *cosa-1(tm3298*) *n*=41, 11.7 ± 0.5; *rmh-1(jf92) n*=61, 7.8 ± 1.3; *zhp-1/2(H25A) n*=163, 6.5 ± 0.8; *zhp-1/2(H25A); zhp-4(vv103) n*=67, 12.2 ± 0.4; *zhp-1/2(H25A); cosa-1(tm3298) n*=42, 11.8 ± 0.6; *zhp-1/2(H25A); rmh-1(vv149) n*=76, 11.5 ± 0.9. Statistical significance assessed by Kruskal-Wallis test and post Dunn’s test: **** *P*<0.0001. (C) Scatterplot graph showing the number of DAPI bodies in the −1 diakinesis oocytes of the indicated genotypes. Error bars denote the mean ± standard deviation. Nuclei scored (*n*), mean ± standard deviation: *+/+ n*=80, *zhp-2(vv84) n*=59, *msh-5(me23) n*=48, *zhp-2(vv84);msh-5(me23) n*=57. Statistical significance assessed by Kruskal-Wallis test and post Dunn’s test: ns=not significant (*P*≥0.05). *msh-5(me23)* to *msh-5(me23);zhp-2(vv84) P*=0.4215. (D, E) Immunofluorescence micrographs of representative nuclei in diplotene, diakinesis, and a magnified bivalent (asterisk) marked with HTP-3 (magenta) and HTP-1 (yellow) of indicated genotypes. Scale bars, 5 µm.

### RMH-1/RMI1 and HIM-6/BLM accumulate at interhomolog HR intermediates in *zhp-2* mutants

Pivotal events of meiotic recombination converge on RMH-1/RMI1, which scaffolds HIM-6/BLM and TOP-3 to form the BTR complex.^23,24,37^ In early pachytene stages, RMH-1 and HIM-6 first colocalize at numerous small foci at RAD-51 processed HR sites, and gradually dissipate from these early HR intermediates while accumulating at others to form ∼6 large late pachytene foci that correspond to designated CO sites (Fig. 3A, E). This dynamic reflects the recruitment of the BTR complex to both CO and nCO intermediates ^23^

**Figure 3.**
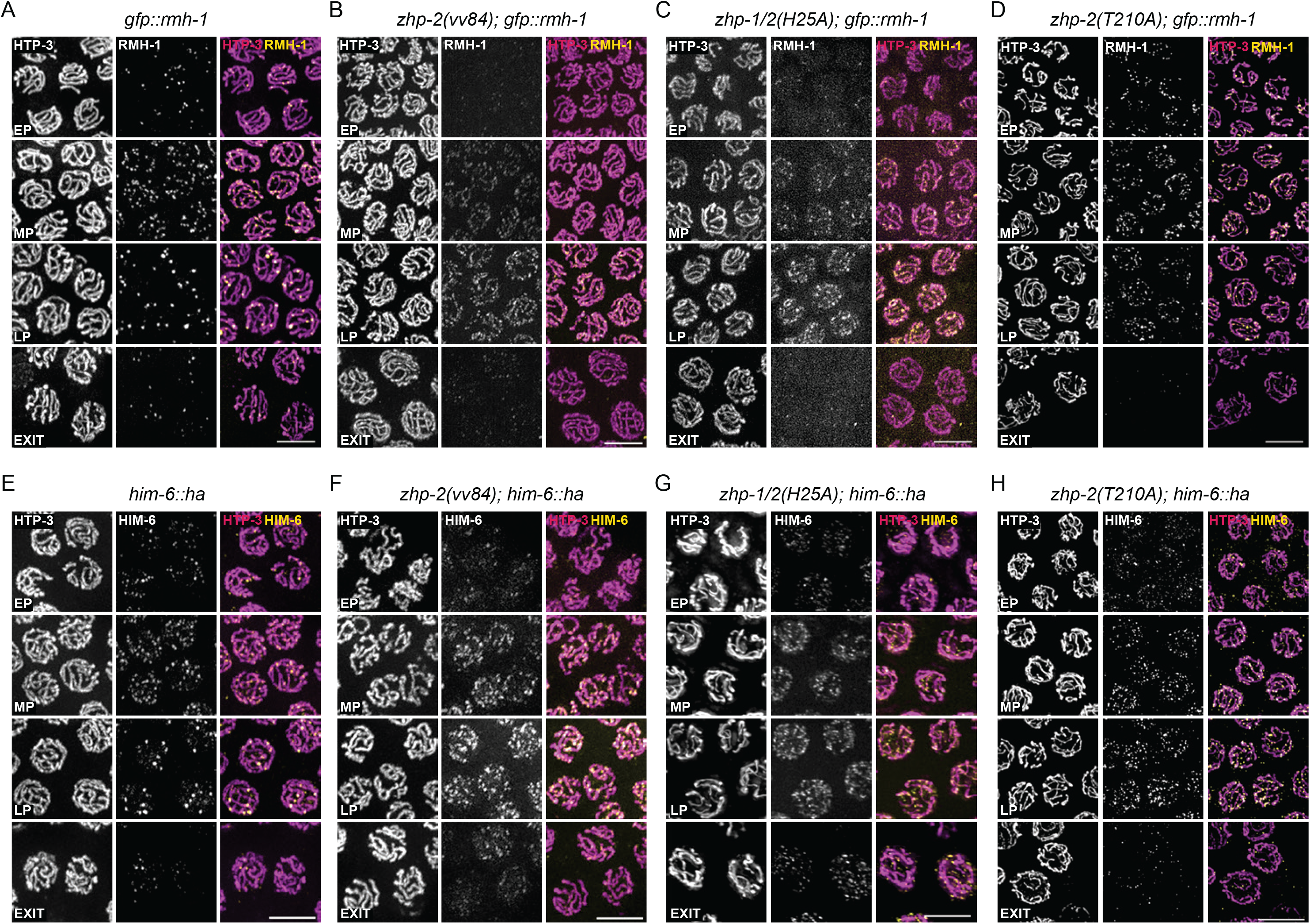
Accumulation of BTR in *zhp*-2 mutants. (A-D) Immunofluorescence micrographs of representative nuclei in early, mid, late pachytene, and pachytene exit (A-D) marked with HTP-3 (magenta) and RMH-1 (yellow) and (E-H) marked with HTP-3 (magenta) and HIM-6 (yellow). Scale bars, 5 µm.

In *zhp-2(vv84*) mutants, small and faint RMH-1 foci first appeared on time in early pachytene nuclei, but with reduced numbers in comparison to wild-type nuclei at the same stage (Fig. 3B). As pachytene progressed however, these foci rapidly increased in size and number, often merging into contiguous stretches that colocalized with the synapsed chromosome axes, and abruptly disappeared at pachytene exit. The hyperaccumulation of RMH-1foci in *zhp-2(vv84)* mutants did not represent ectopic association of the protein with chromosomes outside of the context of HR since RMH-1 foci could not be detected in *spo-11; zhp-2(vv84*) mutants (Fig. S5), indicating that RMH-1 persistence in *zhp-2(vv84)* mutants is a consequence of meiotic recombination initiation.

RMH-1 acts as a scaffold for the BTR complex and is required for localization of HIM-6^23^.To determine if the aberrant RMH-1 localization in *zhp-1/2(vv84)* mutants corresponded to BTR complex dynamics, we examined the localization of HIM-6/BLM, whose appearance and disappearance during wild type pachytene largely mimics the RMH-1 pattern of localization (Fig. 3A, E).^23^ In *zhp-2(vv84*) mutants, HIM-6 repeated the aberrant localization observed with RMH-1; a delayed appearance of a small number of dim HIM-6 foci at early pachytene that rapidly accumulated as pachytene progressed and disappeared at pachytene exit (Fig. 3F). The localization of HIM-6 in mid/late pachytene nuclei also shared important features with the RMH-1 pattern observed in *zhp-2(vv84)* mutants, including foci too numerous to count, and regions in which HIM-6 appeared as stretches that colocalized with the synapsed chromosomes. RMH-1/HIM-6 also accumulated in *zhp-1/2(H25A)* mutants, but at visibly reduced levels in comparison to the null mutant (Fig. 3C, G), consistent with ZHP-1/2 activity at the SC being required for BTR removal from early HR intermediates. Taken together, the observation that HIM-6 reproduces the abnormal localization dynamics of RMH-1 in *zhp-2* mutants is most simply explained by recruitment of the BTR complex to the majority of HR intermediates where it persists in the absence of ZHP-1/2. Given that these intermediates are RAD-51 processed, but fail to convert into COs, the high levels of RMH-1/HIM-6 foci that appear in *zhp-2(vv84)* mutants correspond to sites at which post-strand exchange HR intermediates have accumulated. In summary, our results indicate that ZHP-1/2 association with the SC is required for RMH-1/HIM-6 coarsening; its progressive removal from early HR intermediates and restriction to 6 RMH-1/HIM-6-marked CO sites at late pachytene.

### ZHP-1/2 regulate RMH-1 dynamics at SC polycomplexes

In *C. elegans* mutant backgrounds that disrupt axis morphogenesis, SC components form one or two nuclear aggregates (known as polycomplexes; PCs), a widely observed phenomenon of meiosis under conditions that are not permissive for synapsis.^55^ Nematode PCs mimic aspects of SC organization at the ultrastructural level^56^ and the SC aggregates that form in axis-defective mutants can recruit proCO factors; these show localization dynamics reminiscent of their behaviour during wild-type meiosis.^34,57,58^ For example, ZHP-3 is first found throughout the SC aggregate, but then forms multiple small foci that are eventually replaced by a single large focus containing COSA-1 at late prophase.^34^ To better understand how ZHP-1/2 regulates RMH-1 dynamics during wild-type meiosis, we used the PCs that form in the absence of the axis component HIM-3^10^ to investigate ZHP-2 localization and RMH-1 foci formation. *him-3(gk149)* mutants are competent for wild-type levels of meiotic DSB formation and efficiently use the sister chromatid as a repair template to restore genomic integrity.^10^ We took advantage of this background to first investigate if RMH-1 functions in sister chromatid repair by screening for chromatin-localized RMH-1 foci. In *him-3(gk149)* mutants, PCs first appeared as elongated structures that assumed a compact ball-like form by late pachytene; ZHP-2 colocalized with SYP-1 into the PC, consistent with the SC-dependent localization of ZHP-1/2 observed in wild-type meiosis (Fig. 4A). Importantly, RMH-1 was undetectable on the unsynapsed chromosomes undergoing HR-mediated sister chromatid repair (Fig. 4B). Instead, RMH-1 repeated the behaviour of other pro-CO factors by tracking with SYP-1 into the SC aggregate, providing additional evidence that RMH-1 localization is a specific feature of interhomolog HR. In the majority of these complexes, RMH-1 appeared as a single late prophase focus that disappeared at prophase exit, similar to the dynamic observed during wild-type meiosis. We next investigated the pattern of RMH-1 localization in the PCs when ZHP-2 function is disrupted, with the goal of understanding the mechanism of ZHP-2-mediated regulation of RMH-1. Since we could not recover *zhp-2(vv84*) in combination with *him-3(gk149)*, we turned to *zhp-1/2(H25A)* mutants to probe the effect of reduced ZHP-1/2 activity on RMH-1 localization. In *him-3(gk149); zhp-1/2(H25A)* mutants, ZHP-1/2^H25A^ localized to the PC, but resulted in a dramatic change in RMH-1 localization which resembled the dynamics of RMH-1 in the presence of ZHP-1/2^H25A^ on synapsed chromosomes (Fig. 4B). In PCs containing mutant ZHP-1/2^H25A^, RMH-1 first appeared as numerous heterogeneous foci at the surface of the PC that varied in number, size, and intensity. By late prophase however, RMH-1 foci were reduced to a single bright focus that disappeared at prophase exit, similar to the localization of RMH-1 in the PCs that contained the wild-type ZHP-1/2 complex at this stage. The accumulation and persistence of RMH-1 foci on PCs containing mutant ZHP-1/2^H25A^ complexes, and the hyperaccumulation of RMH-1/HIM-6 in *zhp-2(vv84*) mutants strongly suggests that ZHP-1/2 is a negative regulator of BTR localization, a function that is required for CO formation.

**Figure 4.**
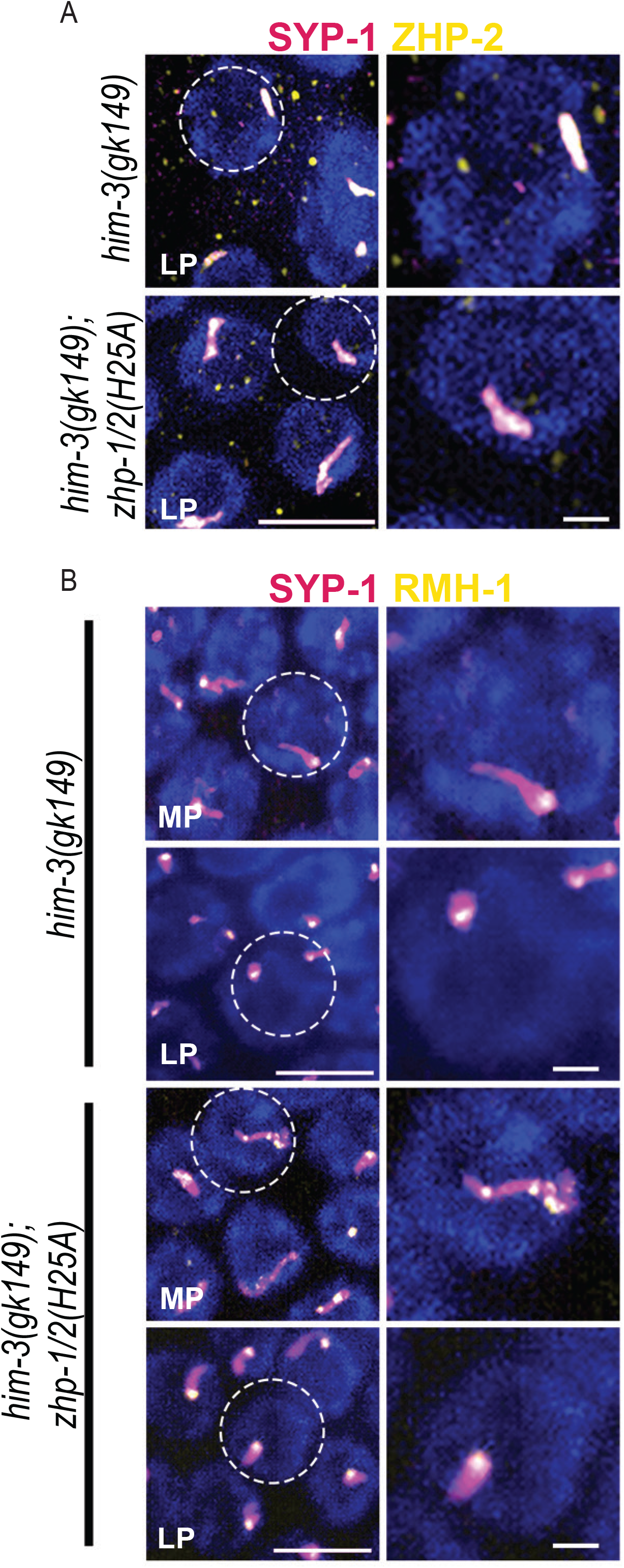
RMH-1 and ZHP-2 localize to SYP-1 aggregates in *him-3(gk149)* mutants. (A-B) Representative immunofluorescence nuclei of mid pachytene (MP) and late pachytene (LP) marked with SYP-1 (magenta), ZHP-2 or GFP::RMH-1 (yellow), and DAPI (blue) from germlines of the indicated genotypes. Scale bars, 5 µm. Magnified single nuclei are encircled. Scale bars, 1 µm.

### ZHP-2 is essential for interhomolog meiotic DSB repair

In *zhp-2(vv84*) mutants, RAD-51 and persisting RMH-1/HIM-6 foci abruptly disappear at late pachytene-pachytene exit stages (Fig. 3B, 5B) and chromosomes emerge intact at diakinesis (Fig. 1D), indicating that meiotic DSBs are efficiently repaired. Although meiotic DSBs can be repaired with a sister chromatid as a template, an interhomolog repair bias dominates until pachytene exit, when sister chromatid repair is licensed to restore genomic integrity.^10,59,60^ In *C. elegans*, the BRC-1/BRD-1 heterodimer (mammalian BRCA1/BARD1) is essential for sister chromatid HR but dispensable during normal meiosis.^59,60^ In *brc-1 brd-1* mutants, RAD-51-marked meiotic recombination intermediates appear and disappear with wild-type kinetics (Fig. 5D) and diakinesis oocytes contain six intact bivalents (Fig. 6A, B), indicating that sister chromatid HR is not a significant pathway of meiotic DSB repair when interhomolog HR is intact. However, when the interhomolog pathway is removed in *brc-1 brd-1; syp-2* mutants that lack SC, the requirement for sister chromatid repair is revealed; diakinesis nuclei containing a spectrum of decondensed chromatin, chromosome fragments, and univalents that are typical of HR-defective *rad-51* mutants.^59,60^ Because chromosomes are fully synapsed in *zhp-2* mutants, both the homolog and the sister chromatid are available as DSB repair templates and we investigated which template is used to repair the persisting late pachytene HR intermediates when ZHP-1/2 is absent. In *zhp-2(vv84)* mutants, the loss of *brc-1 brd-1* resulted in diakinesis nuclei containing masses of decondensed and fragmented chromosomes. The severity of this phenotype mimicked the consequences of loss of RAD-51 itself, indicating the presence of catastrophic levels of unrepaired DSBs (Fig. 6A, B). These results demonstrate that *zhp-2(vv84)* mutants are defective in interhomolog HR and rely on the sister chromatid for DSB repair, revealing that the ZHP-1/2 complex is required for both CO and nCO interhomolog HR. If true, we reasoned that even reduced ZHP-1/2 function should restore some interhomolog HR and mitigate the requirement for sister chromatid-based DSB repair. In *zhp-1/2(H25A)* mutants with reduced levels of the ZHP-1/2 complex at the SC, the loss of *brc-1 brd-1* had no detectable effect on meiotic diakinesis chromosome structure (Fig. 6A, B), indicating that the ZHP-1/2 complex is a potent effector of the interhomolog pathway of meiotic DSB repair.

**Figure 5.**
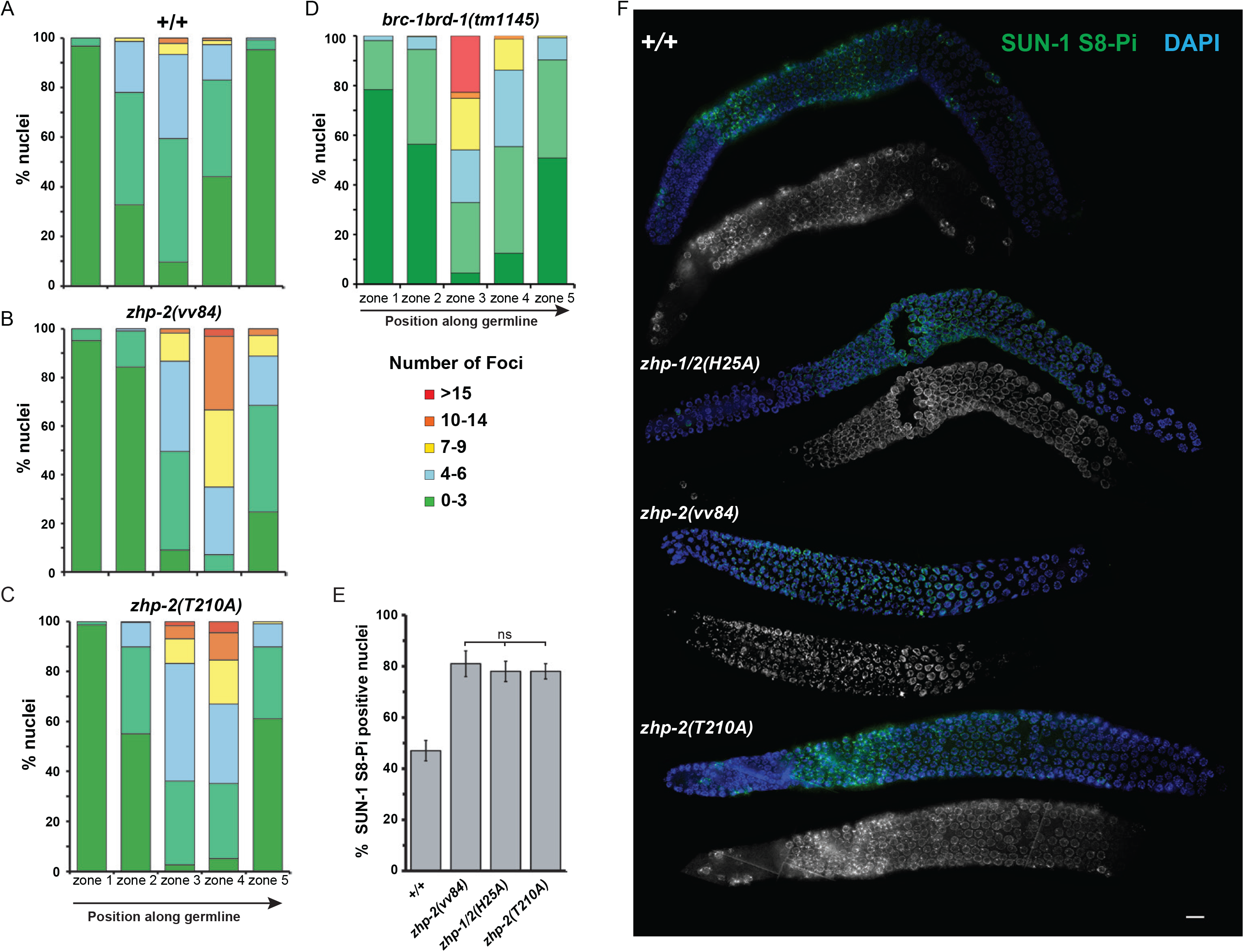
The window of meiotic DSB formation is extended in *zhp-2* mutants. (A-D) Bar graphs showing the percentage of nuclei binned to a range of RAD-51 foci counts per nucleus in respective zones: zone 1 = transition zone, zone 2 = early pachytene, zone 3 = mid pachytene, zone 4 = late pachytene, zone 5 = pachytene exit. Nuclei scored (*n*) in zone 1, 2, 3, 4, 5, percentage of nuclei in zone 1, 2, 3, 4, 5(0-3, 4-6, 7-9, 10-14, >15): *+/+* (383, 247, 303, 251, 250), 1(97.4, 2.6), 2(85.0, 14.6, 0.4), 3(10.9, 58.1, 24.8, 5.9, 0.3), 4(10.0, 57.8, 23.1, 6.8, 1.2), 5(81.6, 17.6, 0.8); *zhp-2(vv84)* (351, 235, 232, 195, 178), 1(95.2, 4.8), 2(84.3, 14.9, 0.9), 3(9.1, 40.5, 37.1, 11.6, 1.7), 4(0.0, 7.2, 27.7, 31.8, 30.3, 3.1), 5(24.7, 43.8, 20.2, 8.4, 2.8); *zhp-2(T210A)* (307, 247, 232, 176, 108), 1(98.7, 1.3), 2(55.1, 34.8, 9.7, 0.4), 3(2.6, 33.6, 47.0, 9.9, 5.2, 1.7), 3(2.6, 33.6, 47.0, 9.9, 5.2, 1.7), 4(5.1, 30.1, 31.8, 17.6, 10.8, 4.5), 5(61.1, 28.7, 9.3, 0.9); *brc-1 brd-1* (314, 330, 376, 305, 248), 1(78.3, 19.7, 1.9), 2(56.4, 38.2, 5.2, 0.3), 3(5.9, 36.7, 27.4, 26.9, 3.2), 4(12.5, 43.0, 30.8, 12.5, 1.3), 5(50.8, 39.5, 8.9, 0.8). (E) Histogram graph showing the percentage of meiotic nuclei positive for SUN-1 S8-Pi in germlines of the indicated genotypes. Nuclei scored (*n*), mean ± standard deviation: *+/+* (1011), 45.6 ± 4.1; *zhp-2(vv84)* (542), 87.11 ± 13.7, *zhp-2(T210A)* (945), 80.9 ± 0.8; *zhp-1/2(H25A)* (750), 84.9 ± 5.1. Statistical significance assessed by Chi-squared test followed by Bonferoni correction: ns=not significant (*P*≥0.05). *zhp-2(vv84)* to *zhp-1/2(H25A) P*= 0.0202; to *zhp-2(T210A) P*= 0.2636 (F) Immunofluorescence micrograph of germlines marked with DAPI (blue) and SUN-1 S8-Pi (green). Scale bar, 10 µm.

**Figure 6.**
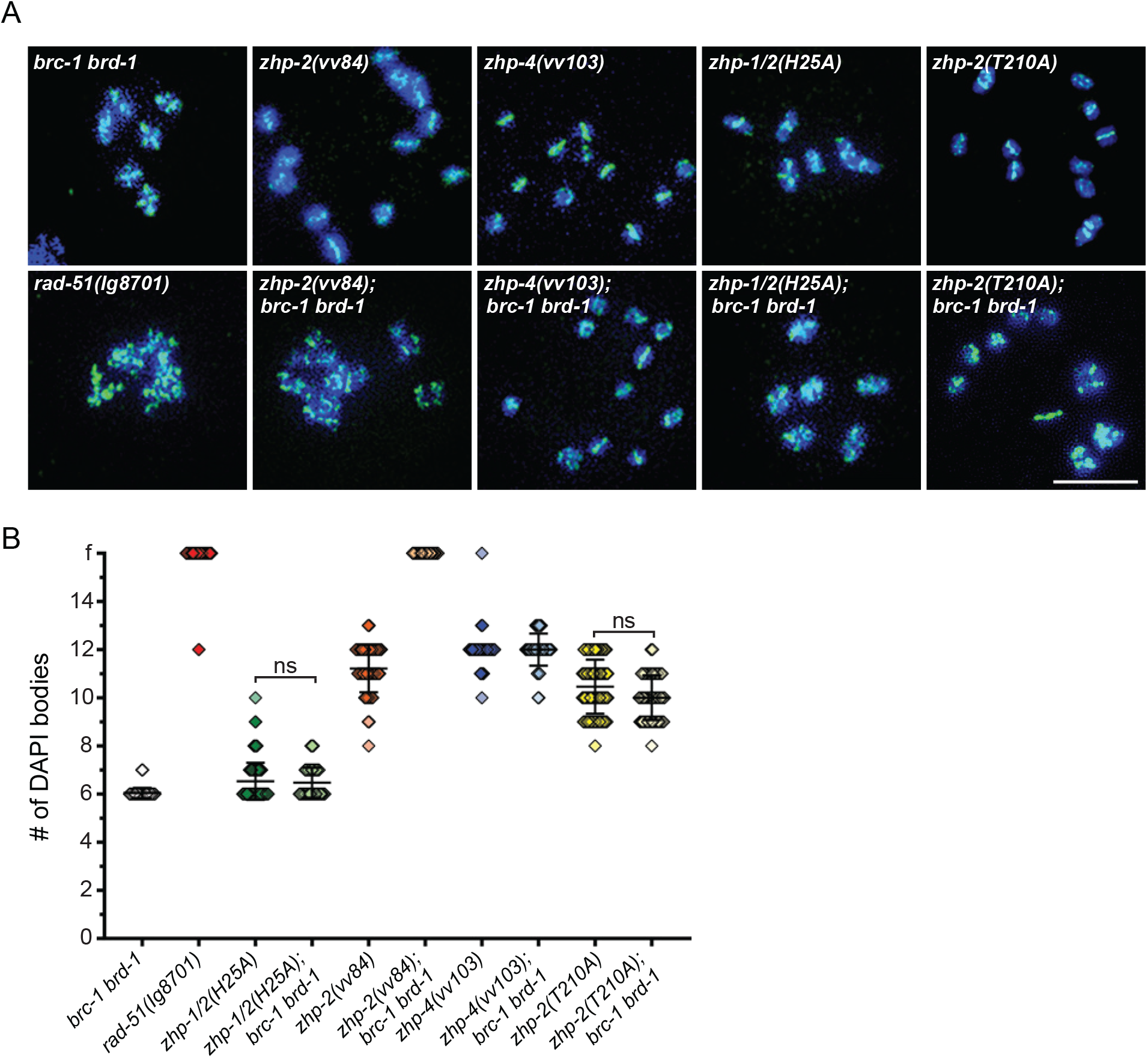
ZHP-2 is required for interhomolog repair of meiotic DSBs. (A) Immunofluorescence micrographs marked with HTP-3 (green) and DAPI (blue) of representative diakinesis −1 oocytes of the indicated genotypes. Scale bars, 5 µm. (B) Scatterplot graph showing the number of DAPI bodies in the −1 diakinesis oocytes of the indicated genotypes. Error bars denote the mean ± standard deviation. Nuclei scored (*n*), mean ± standard deviation: *brc-1 brd-1 n*=85, *rad-51(lg9701) n*=59, *zhp-1/2(H25A) n*=164, *brc-1 brd-1 n*=72, *zhp-2(vv84) n*=56, *zhp-2(vv84); brc-1 brd-1 n*=50, *zhp-4(vv103) n*=79, *zhp-4(vv103); brc-1 brd-1 n*=46, *zhp-2(T210A) n*=100, *zhp-2(T210A); brc-1 brd-1 n*=50. Statistical significance assessed by Kruskal-Wallis test and post Dunn’s test: ns=not significant (*P*≥0.05), **** *P*<0.0001. *brc-1 brd-1* to *zhp-1/2(H25A) P*= 0.0858, to *zhp-1/2(H25A); brc-1 brd-1 P*=0.2032. *zhp-1/2(H25A)* to *zhp-1/2(H25A); brc-1 brd-1 P*= 1. *zhp-2(T210A)* to *zhp-2(T210A); brc-1 brd-1 P*=0.6566.

In *C. elegans*, CHK-2 (checkpoint Kinase-2) regulates many of the early events of meiotic prophase by delaying cell-cycle progression in response to defects in pairing, synapsis, and CO formation.^61–63^ Defects in the establishment of CO precursors results in a CHK-2-mediated extension of the window of DSB induction as a response to activation of the crossover assurance checkpoint; once CO precursors form, CHK-2 is inactivated by PLK-2 (Polo kinase 2) and DSB induction ceases.^63–65^ In *zhp-2(vv84)* mutant germlines, RAD-51-marked recombination intermediates appear at high levels during pachytene, suggesting that the window of DSB induction is extended (Fig. 3B, 5B). To investigate this possibility, we used the phosphorylation of SUN-1 at serine 8 as a proxy to detect CHK-2 activity during meiotic prophase (Fig. 5E, F).^33,66,67^ In wild-type germlines, 47% of prophase nuclei were positive for SUN-1^S8-Pi^, a population that spanned the region from leptotene-zygotene to midpachytene. In *zhp-2(vv84*) mutants, however, 81% of the nuclei were positive for SUN-1 phosphorylation and this population extended into the late pachytene region (*P*<0.0001 in comparison to wild type), revealing a ∼2-fold extension of the window of DSB induction. Taken together, the combined evidence suggests that ZHP-1/2 is required for the transition of interhomolog post-strand exchange HR intermediates into the key CO precursors that can satisfy the crossover assurance checkpoint, trigger CHK-2 inactivation, and end meiotic DSB induction.

### *zhp-2* mutants are defective in CO designation and establishing CO interference

The highly conserved MutSγ complex is required at multiple steps during crossover progression: the stabilization of single end invasions, the formation of joint molecules, and dHJ processing into COs.^68–70^ In *C. elegans,* RMH-1 is required for the appearance of MSH-5 foci, consistent with a role for MSH-5 at early HR intermediates as well as at CO precursors.^37^ MSH-5 is required for the formation of COSA-1 foci, a marker of CO progression,^30^ and COSA-1/CDK-2 are in turn required for accumulation of MSH-5 at COSA-1 foci, revealing a positive feedback loop that progressively concentrates and stabilizes proCO factors at future CO sites.^35^ To characterize the role of ZHP-2 in proCO factor dynamics, we first examined the localization of MSH-5 and COSA-1 in *zhp-2* mutants (Fig. 7A-C). In comparison to the 6 large and bright MSH-5 and COSA-1 foci present in wild-type late pachytene nuclei, highly elevated numbers of smaller foci of varying sizes and intensities appeared in nuclei of *zhp-2(vv84)* mutants at the same stage (average of 39 MSH-5 and 19 COSA-1), similar to the numbers previously reported for *zhp-1-*depleted germlines.^34^ The two-fold elevation of MSH-5 versus COSA-1 foci number is likely to reflect MSH-5 localization to both early and late HR intermediates while COSA-1 is enriched at only a subset of MSH-5 foci that correspond to CO-competent intermediates. These results indicate that ZHP-1/2 is not required for MSH-5 and COSA-1 (or RMH-1/HIM-6) recruitment or stabilization at strand exchange intermediate. Instead, the complex plays an essential role in CO formation by inducing proCO factor accumulation at CO precursors and their disappearance from nCO sites.

**Figure 7.**
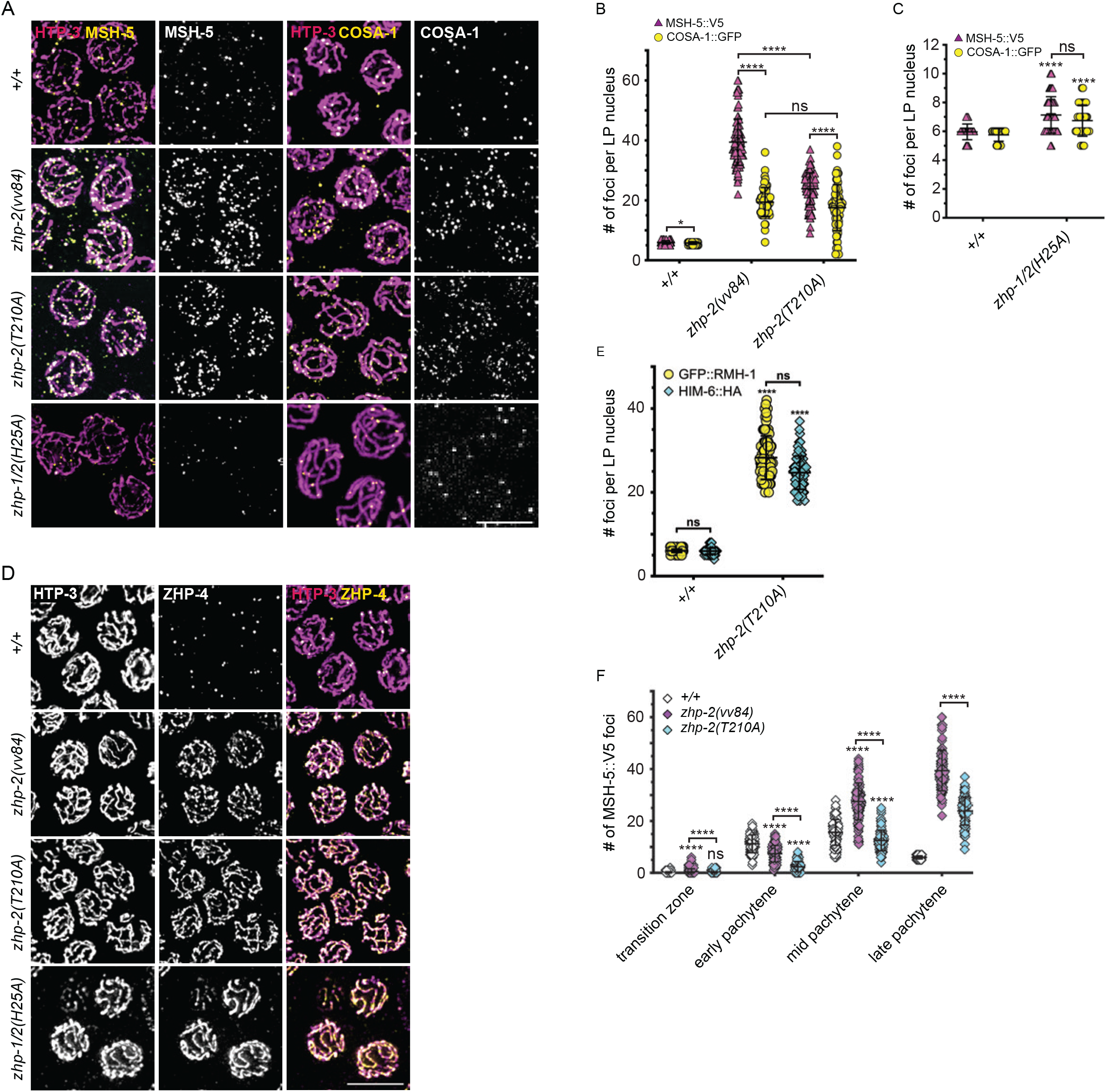
CO formation and designation in *zhp-2* mutants. (A) Immunofluorescence micrographs of representative late pachytene nuclei marked with HTP-3 (magenta) and MSH-5 or COSA-1 (yellow) of indicated genotypes. Scale bar, 5 µm. (B, C) Scatterplot graphs showing the number of MSH-5 and COSA-1 foci in late pachytene nuclei of the indicated genotypes. Error bars denote the mean ± standard deviation. Nuclei scored (*n*) for MSH-5/COSA-1 foci: +/+ *n*=101/64, *zhp-2(vv84) n*=99/60, *zhp-2(T210A*) *n*=101/66, *zhp-1/2(H25A) n*=77/38. Statistical significance assessed by Two-tailed Mann-Whitney tests: ns=not significant (*P*≥0.05), * *P*<0.01, **** *P*<0.0001. MSH-5 to COSA-1 foci of +/+ *P*=0.0136; COSA-1 of *zhp-2(vv84*) to *zhp-2(T210A) P*=0.1335. MSH-5 to COSA-1 of *zhp-1/2(H25A*) *P*= 0.1664. (D) Immunofluorescence micrographs of representative late pachytene nuclei marked with HTP-3 (magenta) and ZHP-4 (yellow) in germlines of the indicated genotypes. Scale bar, 5 µm. (E) Scatterplot graph showing the number of RMH-1 and HIM-6 foci in late pachytene nuclei of the indicated genotypes. Error bars denote the mean ± standard deviation. Nuclei scored (*n*) for RMH-1/HIM-6 foci: *+/*+ *n*=100/95, *zhp-2(T210A)* n= 89/95. Statistical significance assessed by Kruskal-Wallis test and post Dunn’s test: ns=not significant (*P*≥0.05), **** *P*<0.0001. RMH-1 foci to HIM-6: +/+ *P*= 1; *zhp-2(T210A) P*=0.1118. (F) Scatterplot graph showing the number of MSH-5 foci per nucleus of the indicated genotypes at the indicated stages. Nuclei scored (*n*) in transition zone, early, mid, and late pachytene: +/+ (107, 101, 101, 101), *zhp-2(vv84)* (101, 100, 99, 99), *zhp-2(T210A)* (126, 128, 120, 101). Statistical significance assessed by Kruskal-Wallis test and post Dunn’s test: ns=not significant (*P*≥0.05), **** *P*<0.0001. Transition zone *+/+* to *zhp-2(T210A) P*=0.100811.

In contrast to *zhp-2(vv84)* mutants, the late pachytene nuclei of *zhp-1/2(H25A)* RING mutants contained numerous bright MSH-5 and COSA-1 foci that resembled those observed in wild type (Fig. 7A), consistent with the high levels of bivalents observed at diakinesis (Fig. 1E). Unexpectedly, the number of COSA-1 and MSH-5 foci was elevated in comparison to wild-type controls (*P*<0.05; Fig. 7C), suggestive of a disruption in establishing CO interference. To investigate this possibility, three widely spaced SNPs defining a 35 m.u. interval on chromosome V were used to measure the frequency of double-crossing over in oocytes.^52,71^ We calculated the coefficient of coincidence (CoC = 0.303) and derived an interference measure (I = 0.697) that is significantly reduced, indicating that the COs that form in *zhp-1/2(H25A)* mutants are defective in establishing the CO interference that normally limits them to one per homolog pair. To further investigate the functionality of the COs that formed in *zhp-1/2(H25A)* mutants, we assayed their ability to trigger CO-dependent events. We first examined the localization of the ZHP-3/4 complex, which is SC associated until its retraction at late pachytene to 6 bright foci that correspond to CO sites (Fig. 7D).^29,32,34^ In *zhp-2(vv84)* and other CO-defective mutants, ZHP-4 foci do not emerge at late pachytene and instead the complex remains localized with the SC (Fig. 7D).^32,34^ Surprisingly, ZHP-3/4 localization in *zhp-1/2(H25A)* mutants mimicked the dynamics observed in CO-defective mutants, despite the presence of abundant COs. The restriction of ZHP-3/4 to the late pachytene designated CO sites correlates with CO-triggered spatial remodelling of the bivalent that isolates factors required for the loss of sister chromatid cohesion during the meiotic divisions; this includes the retention of the axis component HTP-1 along the long arms,^72,73^ and the restriction of AIR-2/Aurora B kinase to the short arms.^74,75^ In *zhp-1/2(H25A*) mutants, the bivalents failed to establish these CO-triggered changes in chromosome geometry. Instead HTP-1 localized to the axes of both long and short arms while AIR-2 showed strong colocalization with chromatin in addition to its appropriate enrichment at the short arms (Fig. 2D, E, F). These results suggest that the reduced and/or unstable association of ZHP-1/2^H25A^ with the SC can locally support CO designation (as defined by COSA-1 localization) and crossing over, however the COs that form are defective in establishing interference and prompting bivalent remodelling.

### CO formation requires the Polo docking site of ZHP-2

Our data demosntrated that the ZHP-1/2 complex is required for both the removal of proCO factors from HR sites and their stabilization at future CO sites, and a question we sought to address is how ZHP-1/2 triggers proCO factor coarsening. Post-translational modifications of meiotic proteins have previously been shown to be required for CO designation and formation.^35^ The ZHP-1/2 complex has a single S-[pS/pT]-P/X <u>P</u>olo-<u>b</u>ox <u>d</u>omain (PBD) binding motif that is located in the C-terminal intrinsically disordered region (IDR) of ZHP-2, and is predicted to permit Polo kinase binding following the phosphorylation of T210 by a priming kinase.^76^ To determine whether this motif is phosphorylated *in vivo*, we performed a phosphoproteomic analysis using mass spectroscopy of phospho-enriched proteins from *C. elegans* lysates and found that T210 of ZHP-2 was phosphorylated *in vivo* (Fig. S6).^77^ To investigate if mutation of T210 would interfere with polo kinase docking, a T210A substitution was engineered using CRISPR-Cas9 mutagenesis. While *zhp-2(T210A)* mutants showed no defects in pairing, synapsis, or localization of ZHP-2^T210A^ (Fig. 8A, B, S7A, S7B); loss of the PLK docking site resulted in CO defects that were as severe as those of *zhp-2(vv84)* null mutants (Fig. 1D, E), suggesting that Polo kinase recruitment to ZHP-2 is required for the function of the ZHP-1/2 complex in CO formation.

**Figure 8.**
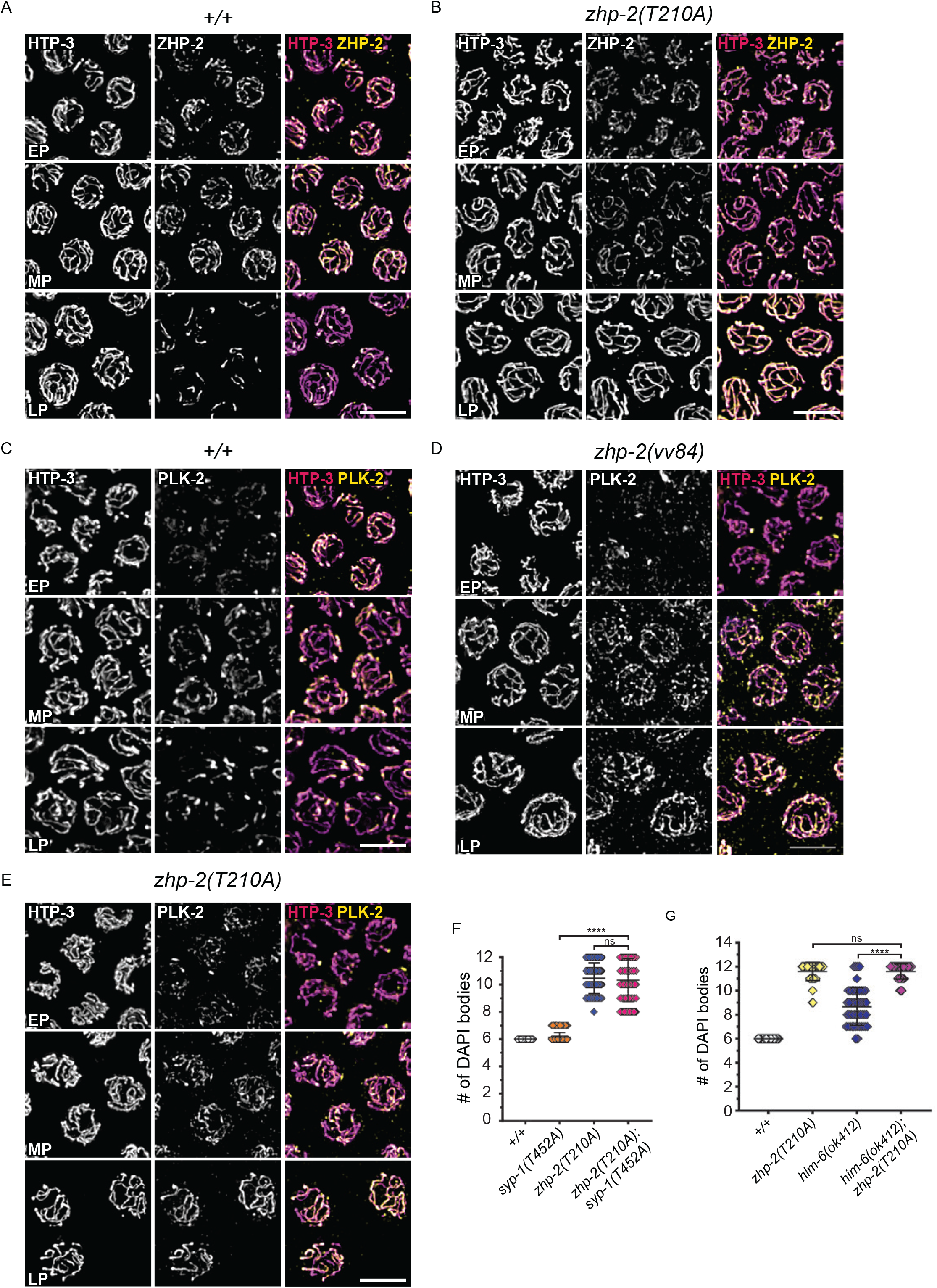
*zhp-2(T210A)* mutants are a separation-of-function allele of *zhp-2*. (A-B) Immunofluorescence micrographs of early (EP), mid (MP), and late pachytene (LP) nuclei marked with HTP-3 (magenta) and ZHP-2 (yellow) from germlines of the indicated genotypes. Scale bars, 5 µm. (C-E) Immunofluorescence micrographs of early (EP), mid (MP), and late pachytene (LP) nuclei marked with HTP-3 (magenta) and PLK-2 (yellow) in germlines of the indicted genotypes. Scale bars, 5 µm. (F) Scatterplot graph showing the number of DAPI bodies in the −1 diakinesis oocytes of the indicated genotypes. Error bars denote the mean ± standard deviation. Nuclei scored (*n*), mean ± standard deviation: *+/+ n*=80, *syp-1(T452A*) *n*=102, *zhp-2(T210A) n*=100, *syp-1(T452A);zhp-2(T210A) n*=78. Statistical significance assessed by Kruskal-Wallis test and post Dunn’s test: ns=not significant (*P*≥0.05), **** *P*<0.0001. *+/+* to *syp-1(T452A) P*= 1, *-2(T210A)* to *syp-1(T452A)zhp-2(T210A) P*= 1. (G) Scatterplot showing the number of DAPI bodies in −1 diakinesis ooctyes of the indicated genotypes. Error bars denote the mean ± standard deviation. Nuclei scored (*n*), mean ± standard deviation: +/+ *n*=46, *zhp-2(T210A) n*=45, *him-6(ok412) n*=51, *him-6(ok412); zhp-2(T210A) n*=48. Statistical significance assessed by Kruskal-Wallis test and post Dunn’s test: ns=not significant (*P*≥0.05), **** *P*<0.0001.

During meiotic prophase, PLK-2 shows a dynamic localization pattern; during leptotene-zygotene it localizes to pairing centers, followed by relocalization to the SC, and then restriction to the short arm of the bivalent in response to CO formation.^67,77–80^ In addition to its localization to chromosome structures, PLK-2 has also been transiently detected at late pachytene CO sites before its rapid relocation to the bivalent short arm.^33,77,80^ Although PLK-2 localized appropriately to the pairing centers and SC in *zhp-2(T210A)* mutants (Fig. 8E), we were unable to determine if its localization was lost at CO sites in *zhp-2(T210A)* germlines given its transient localization and the rare occurrence of COs in these mutants. Instead, we made use of a genetic a background which isolates SC localization of PLK-2 from its localization at CO sites.^63^ In *syp-1(T452A)* mutants, PLK-2 localization to the SC is lost (Fig. S7D), but bivalent formation is unaffected (Fig. 8F),^77^ indicating that any proCO role of PLK-2 occurs independently of the SC-localized population. We reasoned that if CO formation required PLK-2 recruitment to CO sites through containing phosphorylated ZHP-2, the loss of pT210 in *syp-1(T452A)* mutants would lead to a loss in bivalent formation. While *syp-1(T452A)* mutant germlines contained ∼6 bivalents in diakinesis nuclei (Fig. 8F), the number of DAPI-stained structures in diakinesis nuclei of *syp-1(T452A); zhp-2(T210A)* mutants was not different from *zhp-2(T210A)* mutants alone (average of 11, *P*<0.0001), indicating that the PLK-2 docking site on ZHP-2 is required for crossing over in *syp-1(T452A)* mutants. Taken together, these results are consistent with the interpretation that a phosphorylated population of ZHP-2 at the CO site recruits PLK-2, an event which is required for the function of the ZHP-1/2 complex in CO formation.

### ZHP-1/2 localize to CO intermediates on precociously desynapsed chromosomes

To investigate if a CO-localized population of ZHP-2 could be distinguished from the SC-associated population present throughout pachytene, we made use of a phenomenon described in our earlier study^18^; in rare wild-type germlines (<10%), chromosomes spontaneously desynapse at mid-late pachytene, but undergo WT levels of crossing over and bivalent formation by diakinesis.^81^ We used a combination of an endogenous ZHP-1::HA functional fusion tag and ZHP-2 antibodies to investigate the localization of the proteins in germlines showing precociously desynapsed chromosomes as evidenced by the presence of SYP-1 in a nuclear haze or by the presence of separated axes in mid-late pachytene stages.^81^ In wild-type late pachytene nuclei in which SYP-1 appeared nucleoplasmic, ZHP-1 localized to ∼ 6 bright chromosomal foci that disappeared at pachytene exit (Fig. 9C, D). In addition to the ZHP-1 foci in chromosomal regions lacking SYP-1, many nuclei also contained stretches of ZHP-1 that colocalized with stretches of SYP-1 on partially synapsed chromosomes, suggesting that the population of ZHP-1 at the late designated CO sites is qualitatively different and no longer requires the SC to associate with chromosomes. In *zhp-2(T210A)* mutants however, ZHP-2 formed abundant foci in mid-late pachytene that were heterogeneous in size and number, but failed to restrict to 6 foci, mimicking the dynamics of proCO factors in *zhp-2* null mutants (Fig. 9E, F). This suggests that ZHP-2 itself is a proCO factor that functions at HR sites, and that its restriction to the late pachytene CO sites requires polo kinase recruitment to the T210 phosphosite. We suggest that ZHP-1/2 is present in two populations: 1) an SC-associated population that disperses when the chromosomes desynapse, and 2) a subpopulation that is normally concealed by the SC-localized population and is associated with early HR sites, CO intermediates, and late pachytene CO sites. Taken together, these results suggest that ZHP-1/2 functions at HR sites where the complex is required for pro-CO factor coarsening via PLK-2 recruitment, an interpretation that is further supported by our following analysis of proCO factor dynamics in *zhp-2(T210A)* mutants.

**Figure 9.**
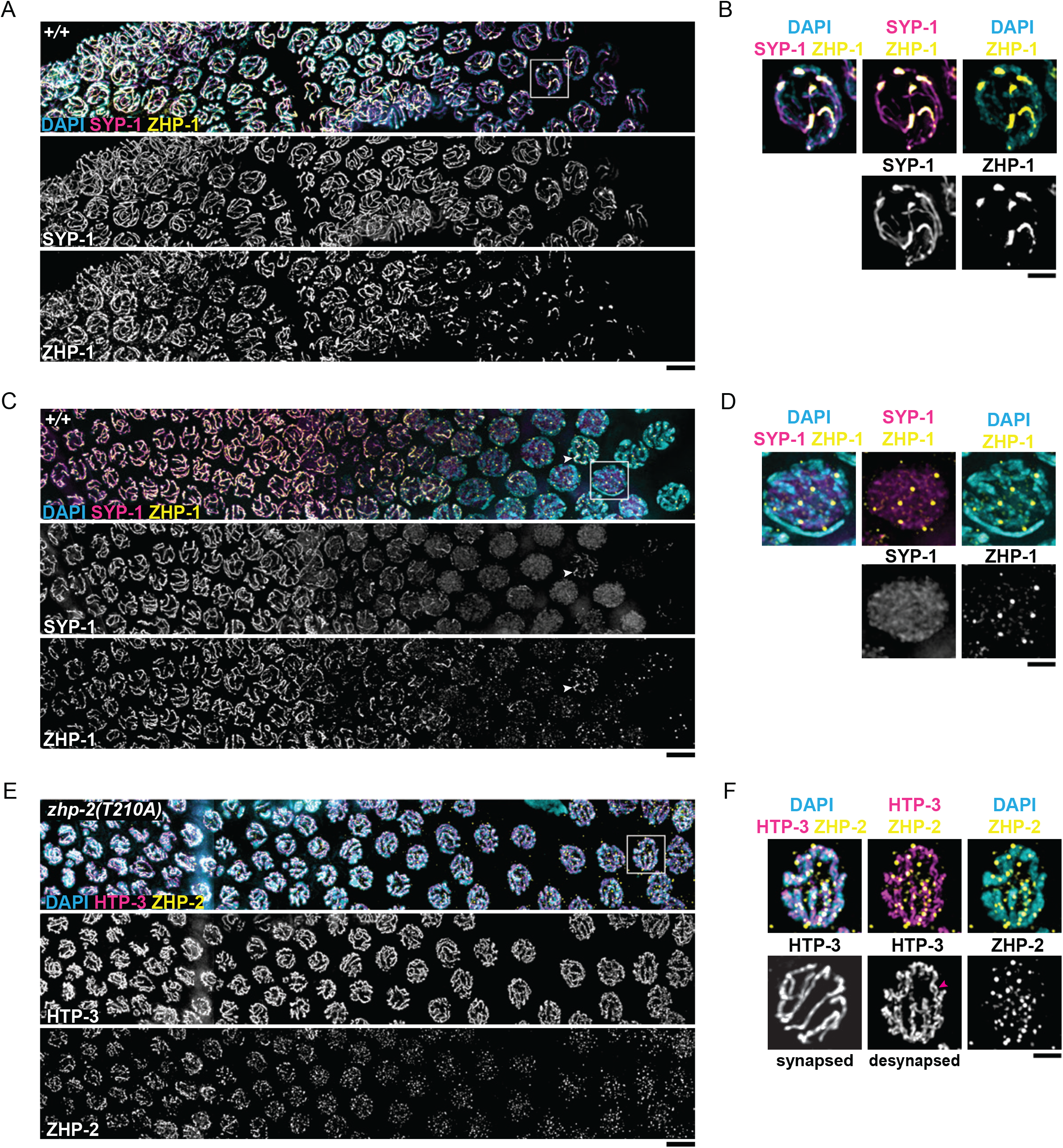
ZHP-1 and ZHP-2 localize as foci in late pachytene on precociously desynapsed chromosomes. (A-F) Immunofluorescence micrographs of germlines marked with SYP-1 (magenta), ZHP-1::HA or ZHP-2 (yellow), and DAPI (blue) in germlines of the indicated genotypes. Scale bars, 5 µm. Enlarged nuclei are denoted by the box. Scale bars, 1 µm.

### Inactivation of the CO assurance checkpoint requires pT210 of ZHP-2

To determine the role of pT210 in CO formation, HR progression in *zhp-2(T210A)* mutants was investigated and compared to the defects observed in *zhp-2(vv84)* mutants in which the ZHP-1/2 complex was not present. As a first step, we examined if the CO assurance checkpoint was satisfied in *zhp-2(T210A)* mutants (Figure 5). The length of the germline region containing SUN-1^S8-Pi^-positive nuclei was not different between *zhp-2(T210A)* and *zhp-2(vv84)* mutants (∼80% for both versus 47% in wild type). Consistent with a prolonged period of DSB induction, *zhp-2(T210A)* mutants also showed elevated levels of RAD-51 foci in comparison to wild type throughout pachytene. These data indicate that the PLK-2 docking site on ZHP-2 is required for the formation of the CO precursor needed for CHK-2 inactivation.

### The Polo docking site of ZHP-2 is required for proCO factor coarsening

To determine how loss of the ZHP-2 phosphosite affects CO progression, we next investigated proCO factor dynamics in *zhp-2(T210A)* mutants. In contrast to the hyperaccumulation of RMH-1/HIM-6 observed in *zhp-2(vv84)* mutants throughout pachytene, RMH-1 and HIM-6 appeared as abundant, but discrete foci in the midpachytene nuclei of *zhp-2(T210A)* mutants, indicating the presence of reduced levels of RMH-1/HIM-6-marked HR intermediates (Fig. 3D, H). At late pachytene, the number of RMH-1 and HIM-6 foci were reduced to similar levels (average of 28 and 25, respectively; *P*=0.1118), but still ∼4.5 times higher than found in wild-type nuclei (Fig. 7E). This suggests that the presence of the ZHP-1/2 complex at the SC *per se* reduces the number of RMH-1/HIM-6-marked intermediates that otherwise persist in the null. This early repair does not require pT210, which is instead required at a later stage to restrict RMH-1/HIM-6 to the designated CO sites at late pachytene. In considering these dynamics and BTR functions, it is probable that ZHP-2 is first required for BTR dynamics at all HR sites (nCO and CO) while the polo docking site on ZHP-2 is required later for restricting and concentrating BTR to the 6 designated CO sites. Recent evidence suggests that HIM-6 is structurally and catalytically required for crossing over by supporting the dHJ conformation and its processing into a CO.^24^ We considered the possibility that the high levels of HIM-6 at late pachytene HR sites in *zhp-2(T210A)* mutants resulted in increased unwinding activity and dissolution of joint molecules, leading to CO failure. *him-6(ok412)* mutants have lower numbers of DAPI figures at diakinesis in comparison to *zhp-2(T210A)* mutants, and to determine if excess HIM-6 activity was responsible for the CO defect in *zhp-2(T210A)* mutants, we investigated if loss of HIM-6 would increase bivalent formation at diakinesis (Fig. 8G). Since the number of DAPI bodies in *him-6(ok412); zhp-2(T210A)* mutants was not different from *zhp-2(T210A)* mutants alone (*P*>0.05), we conclude that the loss of CO formation in the absence of pT210 was independent of HIM-6 activity.

In both *zhp-2(vv84)* and *zhp-2(T210A)* mutants, MSH-5 appeared as discrete foci throughout pachytene, thereby facilitating a time-course analysis of MSH-5 foci dynamics as a marker of CO progression (Fig. 7F). The appearance of MSH-5-marked early recombination intermediates in both the *zhp-2* null and the phosphomutant was initially delayed, but by late pachytene rapidly accumulated to >4-fold higher than the 6 observed in wild type (average of 39 and 24, respectively). Importantly, at midpachytene when MSH-5 marks both CO and nCO sites,^23,33^ the number of MSH-5 foci in wild type and *zhp-2(T210A)* mutants are similar (average of ∼15 and 12, respectively; *P*<0.0001) while the number of foci in *zhp-2(vv84)* mutants are two-fold higher (average of 27). These data collectively suggest that the higher levels of MSH-5 (and RMH-1/HIM-6) foci in *zhp-2(vv84)* mutants in comparison to the phosphomutant are the consequence of persisting early interhomolog HR intermediates that undergo timely repair as nCOs in *zhp-2(T210A)* mutants. If true, we reasoned that CO-defective *zhp-2(T210A)* mutants would be proficient in interhomolog nCO-mediated repair, and we examined the effect of loss of the sister chromatid pathway of HR repair on diakinesis chromosome morphology. In contrast to *zhp-2(vv84); brc-1 brd-1* mutants, the diakinesis nuclei of *zhp-2(T210A); brc-1 brd-1* mutants showed condensed and well-formed chromosomes, indicating that phosphorylation of T210 is not required for nCO repair (Fig. 6A, B). These results strongly suggest ZHP-1/2 is required for both nCO and CO repair of DSBs, while repair as a crossover uniquely requires the PLK-2 docking site on ZHP-2.

In *zhp-2(T210A)* mutants, late pachytene nuclei also showed a ∼3-fold increase in COSA-1 foci in *zhp-2(vv84)* mutants in comparison to wild type, indicating that like MSH-5 and RMH-1/HIM-6, its coarsening and restriction 6 large foci at late pachytene requires ZHP-2 phosphorylation at T210 (Fig. 7B). This suggests that ZHP-1/2 function at the SC can support crossover progression to the 6 designated COs once PLK-2 is recruited to ZHP-2 at the future CO site. Using STED (<u>s</u>timulated <u>e</u>mission <u>d</u>epletion) super-resolution microscopy, structures interpreted as dHJs are visible as a COSA-1 focus flanked by MSH-5 doublets.^24^ Using these criteria, we observed that approximately 15% of the COSA-1 foci in wild-type late pachytene nuclei were in this orientation, while ∼5% were observed in both *zhp-2(vv84)* and *zhp-2(T210A)* mutants (Fig. S8; *P*=0.0003 for both mutants in comparison to wild type). The bias for fewer dHJ-like structures in the mutants can stem from the continuous DSB formation that occurs as part of the CO assurance checkpoint (Fig. 5E, F), and the mixed population of HR intermediates that result. Importantly, these results suggest that *zhp-2* mutants can form dHJ-like intermediates, but fail in executing the last steps required for resolution as COs. an event that requires phosphorylation of the polo docking site. Given the key role of phospho-regulation of proCO factors at HR sites,^35,38,47^ we favor the model that phosphorylation of ZHP-2 at the CO precursor by locally enriched kinases (*i.e.* CDK-2/COSA-1) leads to PLK-2 recruitment to the site. An essential proCO function of PLK-2 is to inactivate and stabilize ZHP-1/2, which in turn stabilizes the proCO assemblies and licenses proCO accumulation, leading to a proCO factor population at the CO site that can support dHJ resolution into a CO. Concomitant with these events, ZHP-1/2 remains active at other HR intermediates and continues to displace RMH-1/HIM-6 and other proCO factors, leading to their dissipation and the repair of the HR site as an nCO.

## Discussion

Our study demonstrates that the ZHP-1/2 complex has both early and late functions in CO formation that are based on regulating the dynamics of proCO factors. In the absence of the complex, these factors accumulate at HR intermediates, but 6 large foci do not emerge at late prophase, and COs fail to form. Thus, ZHP-1/2 negatively regulates proCO factor association with HR intermediates, a process that is required for CO formation. ZHP-1/2-mediated removal of RMH-1/HIM-6 from early HR sites is required for interhomolog repair and CO formation, consistent with a function for the complex in facilitating JM formation through local BTR loss. At later steps in the CO pathway, we demonstrate that phosphorylation of the Polo docking site of ZHP-1/2 at CO precursors triggers coarsening of RMH-1/HIM-6, MSH-5, and COSA-1. Since PLK-2 also functions in the CO assurance pathway by inhibiting CHK-2 in response to CO designation,^63^ our study reveals that ZHP-2 is a central hub that coordinates interhomolog HR with CO formation and inactivation of the crossover assurance checkpoint. Our results strongly suggest that ZHP-2 located at CO precursors is phosphorylated by proximal proCO kinases (*i.e.* CDK-2/COSA-1), a modification that leads to PLK-2 recruitment and initiation of a cascade of events that begin with stabilization of proCO factors at the designated CO site, protection of HJ intermediates, execution of crossing over, and repair of other HR sites as nCOs.

Any model to explain the crossover function of ZHP-1/2 must address the paradox of abundant pro-CO factor-marked recombination intermediates that fail to convert into COs in the absence of the complex. Our results provide strong evidence that ZHP-1/2 are required at two temporally distinct steps in the CO pathway: first in generating the joint molecules required for dHJ formation, and second, in initiating coarsening of proCO factors to convert CO intermediates into COs. Amongst the proCO factors, RMH-1 and HIM-6 have the earliest function in HR and show the most dramatic accumulation in *zhp-2* mutants, followed by a similar failure in forming 6 bright foci at late pachytene that correspond to CO sites. The appearance of similar levels of RMH-1 and HIM-6 suggests that the BTR complex forms and persists, leading to its unscheduled activity at early HR sites. Since *zhp-2(vv84)* null mutants are unable to repair meiotic DSBs using interhomolog CO or nCO pathways, it is reasonable to conclude that BTR persistence disrupts an early step required for all HR-mediated meiotic DSB repair. RMH-1/HIM-6 require RAD-51 processing of DSBs for localization to HR sites;^23^ consequently, the key HR intermediate affected by loss of *zhp-1/2* must have advanced to the post-strand exchange step. BTR complexes have an extensively documented ability to reverse DNA strand invasion intermediates,^82^ and persistence of RMH-1/HIM-6 at these early HR sites in *zhp-2(vv84)* mutants could result in perduring unwinding activity that impedes D-loop formation, a prerequisite for second end capture and dHJ formation, and for repair by SDSA. The possibility that DNA strand exchange intermediates undergo continuous dissolution in *zhp-2(vv84)* mutants predicts that the vast majority of persisting pachytene HR intermediates contain free, processed DNA ends ready to be used in HR-mediated repair. This is supported by the evidence that the levels of unrepaired meiotic DSBs in *zhp-2(vv84)* mutants are extraordinarily high, yet repaired efficiently when the sister chromatid is licenced as a repair template at late pachytene. First, the loss of the late pachytene backup HR pathway in *zhp-2(vv84)* mutants reveals a *rad-51* mutant-like loss genomic integrity at diakinesis, suggestive of comparable levels of unrepaired DSBs. Second, the persisting DSBs in *zhp-2(vv84)* mutants are efficiently repaired with the sister chromatid at pachytene exit, suggesting that the ends have been processed for HR and can be rapidly redirected to the sister chromatid as chromosomes desynapse. Taken together, our results indicate that ZHP-2 is required for dissociation of BTR from HR intermediates, and several lines of evidence suggest that this occurs through a direct interaction: 1) ZHP-1 and ZHP-2 have been found in RMH-1 immunoprecipitates,^37^ 2) the inclusion of ZHP-1/2*^H25A^* in nuclear SC polycomplexes changes RMH-1 dynamics, 3) the presence of even low levels of ZHP-1/2^H25A^ at the SC correlates with a visible reduction in RMH-1/HIM-6 localization.

While the molecular function of ZHP-1/2 is unknown, it is likely to be a ubiquitin E3 ligase given its structural and functional similarity to HEI10. While largely known to target proteins for proteolysis, ubiquitination can also disrupt or disassemble complexes independent of protein degradation.^83^ In the context of crossover formation, such a mechanism is supported by the demonstration that ubiquitination (but not proteasomal degradation) is required for timely dissociation of proCO factors in *C. elegans*, and that human HEI10 catalyzes K63-linked polyubiquitin *in vitro*.^53,84^ These observations suggest a scenario in which ZHP-1/2-catalyzed ubiquitination of proCO factors, leads to destabilization of their interactions at HR sites; however, it is also possible that ZHP-1/2 directly displace or disrupt proCO factors through protein-protein interactions. Our results are compatible with the model that ZHP-1/2 targeting of RMH-1 at strand exchange intermediates results in transient BTR disassembly and creates a window of D-loop stability required for joint molecule formation and second end capture. Thus, an essential early function for ZHP-1/2 in crossing over is in fostering the formation of the early joint molecules that can give rise to CO precursors.

Similar to the aberrant pachytene dynamics displayed by RMH-1 foci in *zhp-2* mutants, MSH-5 and COSA-1-marked foci also persist and accumulate into late pachytene stages and fail to restrict to six designated CO sites. The higher levels of MSH-5 in comparison to COSA-1 is consistent with the earlier role for MSH-5 in stabilizing single end invasions as well as its later role in stabilizing CO-competent intermediates that recruit COSA-1.^30,69^ The fact that *zhp-2(T210)* mutants can form dHJ like-structures (albeit at highly reduced levels), but fail to form COs, suggests that ZHP-1/2 also function at late steps in the HR pathway that include resolution of dHJs into COs. This is supported by the observation that the complex is concentrated at the late pachytene designated CO sites in nuclei that have spontaneously desynapsed, indicating that it is present at the right time and place. The loss of CO formation in *zhp-2* mutants is accompanied by a failure in coarsening of proCO factors, suggesting that the two are mechanistically linked. Since the concentration of proCO factors at the designated CO site stabilizes and protects the dHJ from dissolvases,^24,33^ the loss of proCO factor coarsening in *zhp-2* mutants could render them vulnerable to anti-CO factors. The failure to concentrate COSA-1 at the designated CO site can directly disrupt dHJ resolution since it recruits the machinery required for dHJ resolution into COs.^39^

RMH-1, MSH-5, and COSA-1 do not fully co-localize at recombination foci until their restriction to the 6 CO designated sites in late pachytene, and instead show variably interdependent localization dynamics, suggesting that a critical level of individual proCO factors must accumulate to optimize resolution of a CO-specific intermediate into a CO.^23,33,34,47,80,85^ This scenario also explains the heterogeneous nature of the CO defects observed in *zhp-1/2(H25A)* mutants; while MSH-5 and COSA-1 restrict to the CO sites at late pachytene, RMH-1/HIM-6 do not and this is accompanied by failures in spacing of COs, bivalent remodelling, and restriction of ZHP-3/4 to CO sites. Given that the underlying defect in *zhp-1/2(H25A)* mutants is the reduced or unstable interaction of ZHP-1/2^H25A^ with the chromosomes, the concomitant downstream effects are likely to include changes in the individual levels of proCO factor(s) at late recombination intermediates, which in turn disrupt some CO characteristics while leaving others intact. Consequently, the emergence of a fully “competent” CO intermediate at late prophase is a process that is highly sensitive to the composition of the proCO factor population at the CO site.

Although the coarsening of proCO factors is a widely observed meiotic phenomenon, how the process is triggered remains largely mysterious. In this study, we provide several lines of evidence that ZHP-2-mediated recruitment of PLK-2 to a CO precursor triggers proCO factor coarsening and CO formation. In *zhp-2(T210A)* mutants, the loss of the T210 phosphosite results in the accumulation of RMH-1, MSH-5, and COSA-1 and loss of CO formation, indicating that the PLK-2 docking site is essential for proCO factor coarsening and crossing over. The ZHP-1/2 complex normally colocalizes with the SC throughout prophase, however its localization into ∼6 bright interhomolog foci was revealed in late pachytene nuclei that had undergone precocious desynapsis. Importantly, in *zhp-2(T210A)* mutants lacking the PLK-2 docking site, ZHP-1/2 appeared as numerous persisting foci that resembled the localization pattern of other proCO factors in the *zhp-2* phosphomutants. These results suggest that ZHP-2 coarsening requires PLK-2 recruitment to CO precursors, an observation which is most simply reconciled with the scenario that like ZHP-3/4^47^, ZHP-1/2 diffuses within the SC until it is stabilized at CO precursors. In this case, diffusing ZHP-1/2 destabilizes proCO factor association with HR sites until it is stabilized and inactivated at a CO precursor through PLK-2 recruitment. While PLK-2 may effect the changes in ZHP-1/2 dynamics through phosphorylation, Polo kinases can also cause conformational changes on substrates that are independent of its kinase activity^76^, raising the prospect that the docking of PLK-2 itself could stabilize ZHP-1/2 at the designated CO site. Our study provides evidence that coarsening of ZHP-1/2 and other proCO factors protects CO precursors and concentrates factors required for CO execution at late prophase.

## Acknowledgements

The authors would like to thank the members of the Zetka lab for discussions and critical reading of the manuscript, and Griselda Velez-Aguilera for feedback on parts of the analyses. Antibodies, strains, and other reagents were generously provided by Nicola Silva (Masaryk University), Monica Colaiacovo (Harvard University), and Anne Villeneuve (Stanford University). Nematode strains were also provided by the Caenorhabditis Genetics Center, which is funded by the NIH Office of Research Infrastructure Program (P400D010440). This work was supported by funding from Kakenhi (26K23144, 24K01955) and CREST (JPMJCR24B4) to P.M.C., Kakenhi (25K09508) to A.S-C., the Austrian Science Fund (FWF) (PAT25120231212/Grant-DOI 10.55776/PAT2512023) to V.J., and the Canadian Institutes of Health Research (PJT-173381) to M.Z.

A.C-B and R.D. conceived and designed the analyses, generated strains and reagents, collected the data, performed the analysis, and contributed to writing the paper. S.G., A.S-C, P.M.C., and V.J. contributed analysis tools and data. M.Z. conceived and designed the analyses and wrote the manuscript.

## Experimental Procedures

### Strains and Genetics

Bristol N2 strain was used as the wild-type control in the experiments and cultured at 20°C under standard conditions.^86^ Mutations and rearrangements used in this study are listed in Table S1.

### Measuring Recombination

Recombination was measured in *unc-60 dpy-11/ + + (V)* or *dpy-3 unc-3/ + + (X) cis* heterozygotes controls, or in heterozygotes homozygous for either *zhp-2(vv84)* or *zhp-1/2(H25A).* Recombination frequencies were calculated as previously described in Brenner 1974,^86^ where the frequency (*p*) between two markers was calculated using the formula *p* = 1 - (1 - 2R)^1/2^, where R is the number of visible recombinant individuals divided by the number of total progeny. The total progeny number was calculated as 4/3 X (number of WTs + one recombinant class) to compensate for the inviability of the double homozygote class. Both classes of recombinants were used in the calculation.

To assess DCOs on chromosome V in oocytes, *zhp-1/2(H25A)* or N2 Bristol hermaphrodites were mated with *zhp-1/2(H25A)* mutant males bearing the polymorphisms listed below for 24 hours and then individually plated and assessed for F1 outcross progeny by the presence of ∼50% males.^52^ Young L4 larval stage F1 heterozygous hermaphrodite progeny were picked to individual plates, crossed with N2 males for 24 hours and individually plated. F2 male cross-progeny were scored for SNP markers using the products of single worm PCR and restriction digestions with appropriate endonucleases. SNPs used: pKP5076 (genetic position −17), pKP5097 (genetic position +1), snp_Y17D7B (genetic position +18). Statistical analyses of interference were performed using χ2 tests on 2×2 contingency tables of observed and expected DCOs.^87^

### Antibodies and Immunostaining

Anti-ZHP-2 rabbit antibody was raised using a synthetic peptide (CSNIDYSRRDHRNRL) of the C-terminal end of ZHP-2 (GenScript Corporation, Piscataway, NJ) and purified using activated supports according to the manufacturers’ protocols (Affi-Gel 10, BioRad). Specificity of the antibody was tested by immunostaining germlines of *zhp-2(vv84)* mutants and wild-type N2. For whole germline staining 24-26 h post L4-staged adults were dissected in 1X PBS and fixed for 5 minutes in 1% paraformaldehyde on glass slides. After fixation, the gonads were flash-frozen in liquid nitrogen then placed in methanol for 1 minute at −20°C. Three 15-minutes washes with 1X PBS with 1% Tween-20 (PBS-T) were done, followed by blocking with 1% BSA in PBS-T for 30-60 minutes then incubation with primary antibodies overnight in a wet box at 4°C. The following day, three 15-minutes washes with 1% PBS-T were done, followed by 2-hours incubation at room temperature with secondary antibodies in a wet box. Three 15-minutes washes with 1% PBS-T were done prior to mounting in 1 µg/µL of DAPI in Vectashield Mounting Medium (Vector Labs) or for the STED analysis, mounted in Invitrogen ProLong Glass Antifade Mountant (Catalog number P36980) after adding SYBR^TM^ Gold Nucleic Acid Gel Stain (1:600,000, Catalog number S11494) for 1 minute, then washing in 1% PBST-T for 20 minutes. Nuclear spreading was performed for Fig. 7A: gonads were dissected in 5μl of dissecting solution (0.1% Tween-20 in 35% Hank’s Balanced Salt solution) and incubated at 37°C in 50μl of spreading solution (4% w/v paraformaldehyde, 3.2–3.6% w/v sucrose, 1% v/v lipsol, 1% w/v of sarcosyl in water) for 2 hours, followed by incubation in methanol for 20 minutes at −20°C, 3 x 5-minute washes with 1% PBS-T. Blocking and the steps following were performed as previously described. Primary antibodies used in this study: guinea pig *α*-HTP-3 (1:750), rabbit *α*-HTP-3 (1:1000), rabbit *α*-HTP-1 (1:250), rabbit *α*-ZHP-2 (1:750), rabbit *α*-RAD-51 (1:1000), mouse *α*-GFP (1:200, AbCAM ab290), mouse *α*-GFP (1:500, Roche #11814460001), rabbit *α*-OLLAS (1:1000, GenScript, A01658), guinea pig *α*-SUN-1^S8-Pi^ (1:700 gift from V. Jantsch), goat *α*-SYP-1 (1:1000 gift from M. Colaiacovo), rabbit *α*-SYP-1 (1:1000 gift from Nicola Silva), rat *α*-ZHP-4 (1:250), mouse *α*-HA (1:200 BioLegend, 901513), rabbit *α*-AIR-2 (1:250 gift from M. Colaiacovo), rabbit *α*-HIM-8 (1:200 Novus Biological, 41980002), rabbit *α*-PLK-2 (1:200), mouse *α*-V5 (1:200 Invitrogen, AB_2556564). The following secondary antibodies were used: AlexaFluor 555 goat *α*-guinea pig (Molecular Probes, A21435), AlexaFluor 555 goat *α*-rabbit (Invitrogen, A21429), AlexaFluor 555 donkey *α*-goat (Abcam 150130), AlexaFluor 488 donkey *α*-guinea pig (Jackson ImmunoResearch, AB_2340472), AlexaFluor 488 goat *α*-guinea pig (Invitrogen, A-11073), AlexaFluor 488 donkey *α*-rat (Jackson ImmunoResearch, AB_2340684), AlexaFluor 488 goat *α*-rabbit (Molecular Probes, A11034), and AlexaFluor 488 donkey *α*-mouse (Jackson ImmunoResearch, AB_2341099). All secondary antibodies were diluted 1:1000, except: STAR ORANGE goat *α*-rabbit (1:100, Abberior STORANGE-1002-500UG), and STAR RED goat *α*-mouse (1:100, Abberior STRED-1001-500UG).

### Microscopy and image processing

Images were acquired as stacks of 0.2 μm increments using a 100x objective lens on a DeltaVision Deconvolution system equipped with an Olympus 1×70 microscope or a Leica DMI 6000B inverted microscope equipped with a Quorum WaveFX spinning disc and EM CCD camera, or with a 63x objective lens on a ZEISS Axio Observer (wide-field inverted microscope platform). Image stacks were processed and Z-projected using softWoRx 6.5.2 and ImageJ (Fiji) to obtain 2D images from data collected with the DeltaVision and Leica DMI 6000B microscopes. Data collected with the wide-field microscope was deconvolved and Z-projected by ZEISS Zen (blue edition, version 3.7.97.04000) to make 2D images for presentation of figures. Processed images were then finalized using Adobe Photoshop (version 25.12.0) to adjust levels by decreasing background signal. For each observation at least 20 germlines were considered.

Stimulated emission depletion (STED) microscopy images were acquired using Abberior Instruments STEDYCON with an alpha Plan-Apochromat 100x/1.46 Oil DIC. Images acquired from STED were then segmented using the SYBR Gold channel as a reference using the segmentation module available in Cell-ACDC.^88^ SpotMAX was utilized for detection of GFP::MSH-5 and OLLAS::COSA-1 foci.^89^ Parameters were kept a constant for all the analyses performed on the genotypes. Only GFP::RMH-1 within a 150nm radius of OLLAS::COSA-1 were counted. GFP::MSH-5 foci within such a radius were then classified as doublets or triplets if 2 or 3 foci were detected respectively. In the mutants, nuclei with at least 18 OLLAS::COSA-1 foci were used for the analysis presented and in the wild type nuclei with 6 OLLAS::COSA-1 foci in confocal mode. The OLLAS::COSA-1 foci in the mutants are dimmer than in the wild type. Therefore, the mutants were subjected to a 561nm laser excitation power of 34% while the wild type was subjected to 10%.

### Scoring embryonic lethality and incidence of males

L4-stage hermaphrodites were individually plated and transferred to different plates twice a day for three days; on the final brood, the hermaphrodite was left on the plate for 14-16 hours and then removed. The number of eggs on each plate was counted and the plates incubated for 3 days to allow development to adulthood. The number of adult worms were counted, as well as the number of males in the population. Embryonic lethality was determined as the total number of adult worms divided by the total number of eggs laid and the frequency of male progeny was determined as the total number of males divided by the total number of worms.

### Scoring fragmented DAPI bodies

Young adult hermaphrodites were dissected and immunostained as described earlier. The number of DAPI bodies in –1 oocytes (the oocyte next to the spermatheca) were assessed for fragmentation and counted. If DAPI bodies were resolvable individual entities, the number of bodies were counted and presented as the number of DAPI-stained bodies. If the oocyte contained masses of DAPI-stained chromatin that could not be individually resolved, the oocyte was scored as “f” indicating fusion/fragmentation of the chromosomes.

### Scoring foci in staged nuclei

Germline nuclei were staged as follows: mitotic region as nuclei from the distal tip to the row containing condensed crescent shaped nuclei; transition zone as polarized, crescent shaped nuclei; early pachytene as crescent shaped nuclei with some chromatin outside of the crescent; mid pachytene as nuclei containing well-defined tracks dispersed evenly throughout the nucleus; late pachytene as nuclei within the last 5-7 rows before pachytene exit where the chromatin begins to remodel. After immunostaining as described above for RAD-51, RMH-1, HIM-6, MSH-5, and COSA-1, the number of foci colocalized with the axial marker HTP-3 were scored per nucleus in each staged nucleus.

### CRISPR-cas9 genome editing

In an EMS-based screen for recessive mutations that disrupt meiotic chromosome segregation, we isolated a *vv84*, G to A substitution in *zhp-2* that introduces a premature stop codon at the third amino acid (W3STOP) in the predicted protein sequence of *zhp-2.* All other mutants were generated using CRISPR-cas9 genome editing.^49^ CRISPR-cas9 reagents were prepared by mixing 1.25µL of 8µg/µL tracrRNA, 0.3 µL of 8 µg/µL *dpy-10* crRNA, 0.275 µL of 1 µg/µL *dpy-10* template, 0.5 µL of 8 µg/µL target crRNA, 1.1 µL of 1 µg/µL target template, 0.25 µL of 1M KCE, 0.375 µL of 200 µM HEPES pH 7.4, and 6.25 µL cas9. Reagents were centrifuged at 13,000 rpm for 3 minutes and loaded into a glass needle. 24 hours post L4 worms were injected with the reagent in the proximal 1/3^rd^ of one or both gonad arms and allowed to recover overnight at 20°C. Dumpy positive progeny from injected hermaphrodites were isolated and tested for the desired mutation by PCR followed by restriction digest and confirmed by Sanger sequencing. *zhp-1(Δ)* is an indel mutation; a large deletion of the majority of the protein coding sequence (a.a. 12-705 inclusive), resulting in a frameshift and a premature STOP codon. The resulting mutant DNA sequence is ATGGAGTTCAT-TCAGATGTAG. *zhp-1(H25A)* and *zhp-2(H25A)* were created introducing a CA to GC nucleotide substitution (nucleotide 74 and 75), that changed the 25^th^ codon from CAC to GCC. The *zhp-2(T210A)* mutant was created by substituting A (nucleotides 628) to A resulting in a codon change from ACT to GCT. Similarly, *rmh-1(vv149)* was created by introducing a point mutation at nucleotide 35 that replaced G with an A, resulting in a premature stop codon.

### Phosphoproteomics

Mass spectrometry was performed as described in Sato-Carlton *et al.*:^77^ 2 mL pelleted, frozen adult worms harvested in M9 were thawed and dissolved in 5 ml urea lysis buffer (20 mM Hepes, pH 8.0, 9 M urea, 1 mM sodium orthovanadate, 2.5 mM sodium pyrophosphate, and 1 mM β-glycerol-phosphate), sonicated for 1 min at 30-s intervals 10 times, spun down at 20,000 g for 15 min; the supernatant was subjected to PTMScan analysis (Cell Signaling Technology): phosphorylated peptides were enriched by phospho-(Ser/Thr) kinase substrate antibody-immobilized protein A beads and analyzed by liquid chromatography–tandem mass spectrometry using an LTQ-Orbitrap-Elite ESI-CID (Thermo Fisher).

### Statistics and reproducibility

Statistical analyses were performed with Prism 8 (GraphPad) and Microsoft Excel. Embryonic lethality, high incidence of males, and % nuclei positive for SUN-1 S8-Pi were statistically tested using Chi-squared test and Bonferroni corrections (*P* < 0.05 was considered as significant, CI = 0.05: ns (not significant) if *P* > 0.05, \**P* < 0.05, \*\**P* < 0.01, \*\*\**P* < 0.001, \*\*\*\**P* < 0.0001). Distributions of DAPI bodies were statistically tested using Kruskal-Wallis test followed by Dunn’s multiple pairwise comparison tests (*P* < 0.05 was considered as significant, CI = 0.05: ns (not significant) if *P* > 0.05, \**P* < 0.05, \*\**P* < 0.01, \*\*\**P* < 0.001, \*\*\*\**P* < 0.0001). MSH-5:V5, GFP::COSA-1, RMH-1::GFP, HIM-6::HA, and RAD-51 foci scores were statistically tested using Mann-Whitney U test (*P* < 0.05 was considered as significant, CI = 0.05: ns (not significant) if *P* > 0.05, \**P* < 0.05, \*\**P* < 0.01, \*\*\**P* < 0.001, \*\*\*\**P* < 0.0001). No data were excluded from the analyses. The experiments were not randomized, and the investigator were not blinded to allocation during experiments and outcome assessment.

