## Supplemental Information for "ZHP-2/HEI10 generates CO precursors and triggers procrossover factor coarsening and CO formation through Polo kinase recruitment"

Centonza-Brenie *et al.* 2026

### Supplemental Data

Tables S1 and S2

Figures S1-S9

**Table S1. Strain list**

| STRAIN | Genotype | SOURCE |
|---|---|---|
| N2 | Wild type Bristol isolate | CGC |
| EZ404 | *zhp-1(vv111)/hT2 [bli-4(e937) let-?(q782) qIs48] (I;III)* | This study |
| EZ343 | *zhp-2(vv84)/hT2 [bli-4(e937) let-?(q782) qIs48] (I;III)* | This study |
| EZ428 | *zhp-1(vv126[H25A]) I* | This study |
| EZ434 | *zhp-2(vv132[H25A]) I* | This study |
| EZ456 | *zhp-1(vv126[H25A]) zhp-2(vv132[H25A]) I* | This study |
| EZ549 | *zhp-2(vv165[T210A])/tmC18 I* | This study |
| AV106 | *spo-11(ok79)/nT1[unc-?(n754) let-?] (IV;V)* | CGC |
| EZ512 | *zhp-1(vv126[H25A]) zhp-2(vv132[H25A]) I; spo-11(ok79)/nT1[unc-?(n754) let-?] (IV;V)* | This study |
| NSV129 | *msh-5(ddr22[GFP::msh-5]) IV* | Silva Lab |
| EZ511 | *zhp-2(vv84)/hT2 [bli-4(e937) let-?(q782) qIs48] (I;III); msh-5(ddr22[GFP::msh-5]) IV* | This study |
| YMK154 | *msh-5(kim35[msh-5::V5]) IV* | Kim Lab |
| EZ616 | *zhp-2(vv84)/tmC18 I; msh-5(kim35[msh-5::V5]) IV* | This study |
| EZ559 | *zhp-2(vv165[T210A])/tmC18 I; msh-5(kim35[msh-5::V5]) IV* | This study |
| EZ601 | *zhp-1(vv126[H25A]) zhp-2(vv132[H25A]) I; msh-5(kim35[msh-5::V5]) IV* | This study |
| AV630 | *meIs8 [Ppie-1::gfp::cosa-1 + unc-119(+)] II* | CGC |
| EZ353 | *zhp-2(vv84)/hT2 [bli-4(e937) let-?(q782) qIs48] (I;III); meIs8 [Ppie-1::gfp::cosa-1 + unc-119(+)] II* | This study |
| EZ568 | *zhp-2(vv165[T210A])/hT2 [bli-4(e937) let-?(q782) qIs48] (I;III); meIs8 [Ppie-1::gfp::cosa-1 + unc-119(+)] II* | This study |
| EZ72 | *dpy-3(e27) unc-3(e151) X* | This study |
| EZ362 | *zhp-2(vv84)/hT2 [bli-4(e937) let-?(q782) qIs48] (I;III); dpy-3(e27) unc-3(e151) X* | This study |
| EZ513 | *zhp-1(vv126[H25A])zhp-2(vv132[H25A]) I; dpy-3(e27) unc-3(e151) X* | This study |
| DR181 | *unc-60(m35)dpy-11(e224) V* | CGC |
| EZ361 | *zhp-2(vv84)/hT2 [bli-4(e937) let-?(q782) qIs48] (I;III); unc-60(m35)dpy-11(e224) V* | This study |
| EZ514 | *zhp-1(vv126[H25A]) zhp-2(vv132[H25A]) I; unc-60(m35) dpy-11(e224) V* | This study |
| EZ471 | *zhp-1(vv126[H25A]) zhp-2(vv132[H25A]) I; meIs8 [Ppie-1::gfp::cosa-1 + unc-119(+)] II* | This study |
| AV596 | *cosa-1(tm3298)/qC1[qIs26] III* | Yokoo *et al.* 2012 |
| EZ403 | *zhp-4(vv103) V* | Nguyen *et al*. 2018 |

| | | |
|---|---|---|
| UV279 | *rmh-1(jf92[MO1E11.3::unc-119+]) I* | Jagut *et al.* 2016 |
| EZ475 | *zhp-1(vv126[H25A]) zhp-2(vv132[H25A]) I; cosa-1(tm3298)/qC1[qIs26] III* | This study |
| EZ499 | *zhp-1(vv126[H25A]) zhp-2(vv132[H25A]) I; zhp-4(vv103) V* | This study |
| EZ478 | *zhp-1(vv126[H25A]) zhp-2(vv132[H25A]) rmh-1(vv149[W12STOP]) I* | This study |
| VJ1263 | *rmh-1(jf92[M01E11.3::unc-119+]) I; jfSi38 [gfp::rmh-1 cb-unc-119+] II* | Jagut et al. 2016 |
| EZ504 | *zhp-2(vv84)/hT2 [bli-4(e937) let-?(q782) qIs48] (I;III); jfSi38 [gfp::rmh-1 cb-unc-119+] II* | This study |
| EZ491 | *zhp-1(vv126[H25A]) zhp-2(vv132[H25A]) rmh-1(vv149[W12STOP]) I; jfSi38 [gfp::rmh-1 cb-unc-119+] II* | This study |
| EZ662 | *zhp-2(vv165[T210A]) rmh-1(vv149[W12STOP]) I; jfSi38 [gfp::rmh-1 cb-unc-119+] II* | This study |
| EZ514 | *zhp-2(vv84)/hT2 [bli-4(e937) let-?(q782) qIs48] (I;III); jfSi38 [gfp::rmh-1 cb-unc-119+] II; spo-11(me44)/nT1[unc-?(n754) let-?] (IV;V)* | This study |
| UV213 | *him-6(jf93[him-6 ::HA]) IV* | Jagut *et al.* 2016 |
| EZ515 | *zhp-2(vv84)/hT2 [bli-4(e937) let-?(q782) qIs48] (I;III); him-6(jf93[him-6 ::HA]) IV* | This study |
| EZ621 | *zhp-1(vv126[H25A]) zhp-2(vv132[H25A]) I; him-6(jf93[him-6 ::HA]) IV* | This study |
| EZ557 | *zhp-2(vv165[T210A])/tmC18 I; him-6(jf93[him-6 ::HA]) IV* | This study |
| VC418 | *him-3(gk149)/nT1[qIs51] (IV;V)* | CGC |
| EZ490 | *zhp-1(vv126[H25A]) zhp-2(vv132[H25A]) I; jfSi38 [gfp::rmh-1 cb-unc-119+] II; him-3(gk149) IV* | This study |
| EZ489 | *jfSi38 [gfp::rmh-1 cb-unc-119+] II; him-3(gk149) IV* | This study |
| DW102 | *brc-1 brd-1(tm1145) III* | CGC |
| EZ358 | *zhp-2(vv84)/hT2 [bli-4(e937) let-?(q782) qIs48] (I;III); brc-1 brd-1(tm1145) III* | This study |
| CA538 | *rad-51(lg8701)/mIs11 IV* | CGC |
| EZ516 | *brc-1brd-1(tm1145) III; zhp-4(vv103) V* | This study |
| EZ517 | *zhp-1(vv126[H25A]) zhp-2(vv132[H25A]) I; brc-1brd-1(tm1145) III* | This study |
| EZ560 | *zhp-2(vv165[T210A])/hT2 [bli-4(e937) let-?(q782) qIs48] (I;III); brc-1 brd-1(tm1145) III* | This study |
| EZ420 | *zhp-1(vv120[zhp-1:HA]) I* | This study |
| EZ426 | *zhp-1(vv120[zhp-1:HA]) zhp-2(vv84)/hT2 [bli-4(e937) let-?(q782) qIs48] (I;III)* | This study |

| | | |
|---|---|---|
| SSM473 | *iowSi8[pie-1p::gfp1-10::him-3 3UTR + Cbr-unc119(+)] rpa-1(iow89[gfp11::rpa-1]) II* | CGC |
| EZ518 | *zhp-2(vv84)/hT2 [bli-4(e937) let-?(q782) qIs48] (I;III); iowSi8[pie-1p::gfp1-10::him-3 3UTR + Cbr-unc119(+)] rpa-1(iow89[gfp11::rpa-1]) II* | This study |
| EZ502 | *zhp-1(vv126[H25A]) zhp-2(vv132[H25A]) I; iowSi8[pie-1p::gfp1-10::him-3 3UTR + Cbr-unc119(+)] rpa-1(iow89[gfp11::rpa-1]) II* | This study |
| AV307 | *syp-1(me17)/nT1 [unc-?(n754)let-?(qIs50)] (IV;V)* | CGC |
| EZ506 | *jfSi38 [gfp::rmh-1 cb-unc-119+] II; syp-1(me17) V* | This study |
| AV115 | *msh-5(me23)/nT1[unc-?(n754) let-?] (IV;V)* | CGC |
| EZ661 | *zhp-2(vv84) I; msh-5(me23)/nT1[unc-?(n754) let-?] (IV;V)* | This study |
| YKM17 | *plk-2(kim26[plk-2::mRuby]) I; syp-1(kim25[T452A]) V* | Brandt *et al.* 2020 |
| EZ660 | *zhp-2(vv165[T210A]) I; syp-1(kim25[T452A]) V* | This study |
| UV292 | *him-6(ok412) gfp::msh-5 IV* | Jagut *et al.* 2016 |
| UV306 | *zhp-2(T210A)/tmC18 I; gfp::msh-5 IV* | This study |
| UV304 | *zhp-2(T210A)/tmC18 I; him-6(ok412) gfp::msh-5 IV* | This study |
| EZ668 | *ollas::cosa-1 III; gfp::msh-5 IV* | This study |
| EZ656 | *zhp-2(T210A)/tmC18 I; ollas::cosa-1 III; gfp::msh-5 IV* | This study |
| EZ655 | *zhp-2(vv84)/tmC18 I; ollas::cosa-1 III; gfp::msh-5 IV* | This study |

**Table S2. CRISPR materials used in this study**

| Allele | Guide and DNA template | Genotyping primers and detection RE |
|---|---|---|
| *zhp-1(vv111) I* | 5'-ATGGAGTTCATCGTTTGCAA-3' | F: GCTCCCACGTCTTCTGTGAA<br>R: GGTGTCACGTTTTGCTCTGG |
| *zhp-1(vv126[H25A]) I* | 5'-ACAGAAGACGTGGGAGCAGG-3'<br><br>5'-GCAATGGATGTGGATGCTCGCCAAGTAAAAGACAATTTTTTATCACTGCCTGCTCCGCCGTCTTCTGTGAAACATGCCGAACAACTCCCACTGCTGACTTTTGTCATTTGTGTAA-3' | F: GGCACTATGGAGTTCATCGT<br>R: ACATTCTTCGGCAGACTTGC<br><br>BmgBI |
| *zhp-2(vv132[H25A]) I* | 5'-AAACGTGACCGCATGCTGTT-3'<br><br>5'-GCAATCATTGTGGCATTAAACCAAATCAAACGAAACTGTACTTGACAGCATGCGGTGCCGTTTTCTGTCAAAACTGTACGAAAGCTGGTTAGTAAATTTTAATAATAAATTTAGC-3' | F: GGATTGGATCCAATGCAATC<br>R: GTAGCACTGCGAGTTAAATG<br><br>BceAI |
| *zhp-2[vv165(T210A)] I* | 5'-AGGAGTGCTGGTGAGAATCT-3'<br><br>5'-AAGCTAGTCATGAATTTTCATTAAACCTGCGTGATCAGTCAACGTCTAAGATTCTCACCAGCGCTCCTCTCTCCAATGTTAGTTTTATCTCCTGTTTTCCAGTCCCCAGTACACT-3' | F: GGCCAGGCTGAAGGAACTAC<br>R: TTCTTGGCAATACACCCGGC<br><br>HaeII |
| *zhp-1(vv120[zhp-1:HA]) I* | 5'-TCGTCTCAATCGAATCGTGGAGG-3'<br><br>5'-GATCTTCTAGGATTGAGGAATCGATCTGATTCATCATCATCCAACTGCTCGTCTCAAagtAAcaGaGGAGGATCTCTGTTTTACCCATATGATGTCCCGGATTACGCTTAACATAATCTGTAATATAGATGACCATAGTTTTATTTTTCAGCTATAAAC-3' | F: ACTTGGAGGAGGAGGAGGAT<br>R: TGACACACGATTCATTTGGCA |
| *rmh-1(vv149[W12STOP]) I* | 5'-AACTTGATCGTCTTTTCTCT-3'<br><br>5'-CGCATATAAAAACTACAAAATATATGAAAGAAACTGAACTTGATCGTCTTTTCTCTTGACTTGCTAGGAAACATTACCC | F: GTAACCAGCGTACCTGCACA<br>R: CGTTGGGATTTTCATGGTGGG |

| | | |
|---|---|---|
| | ATTCAAGAGAGAATGGCTAAGAATAT GCGTCCAGT-3’ | Hpy188iii |

#### Supplementary Figure Legends

**Figure S1. ZHP-1 and ZHP-2 show codependent colocalization to the SC**
(A) Immunofluorescence micrographs of representative nuclei from mid and late pachytene marked with ZHP-1::HA (green) and ZHP-2 (red) from germlines of the indicated genotypes. (B) Immunofluorescence micrographs of representative nuclei from mid and late pachytene marked with ZHP-2 (green) and HTP-3 (red) from germlines of the indicated genotypes. Scale bars, 5 μm.

**Figure S2. *zhp-1* and *zhp-2* null mutants are proficient for pairing and synapsis**
(A) Immunofluorescence micrographs of representative nuclei from early pachytene marked with HIM-8 (green) and HTP-3 (red) from germlines of the indicated genotypes. Scale bars, 5 μm. (B) Immunofluorescence micrographs of representative pachytene nuclei marked with SYP-1 (green) and HTP-3 (red) from germlines of the indicated genotypes. Scale bars, 5 μm.

**Figure S3. ZHP-2 colocalizes with SC components throughout prophase**
(A) Immunofluorescence micrographs of wild-type nuclei marked with ZHP-2 (red) and HTP-3 (green) of the indicated prophase stages. Scale bars, 5 μm.

**Figure S4. A single intact RING domain is sufficient for localization of the ZHP-1/2 complex**
Immunofluorescence micrographs of representative pachytene nuclei marked with ZHP-2 (green) and HTP-3 (red) from germlines of the indicated genotypes. Scale bars, 5 μm

**Figure S5. RMH-1 accumulation in *zhp-2* mutant depends on the initiation of HR**
Immunofluorescence micrographs of *zhp-2(vv84); spo-11(me44)* mutant nuclei marked with HTP-3 (red) and RMH-1::GFP (green) of the indicated prophase stages. Scale bars, 5 μm.

**Figure S6. T210 of ZHP-2 is phosphorylated**
CID-MS/MS identification of a phosphorylated ZHP-2 peptide. Two MS/MS spectra identified the singly phosphorylated ZHP-2 peptide ILTSTPLSNIDYSR. Both localized the modification to the adjacent S4/T5 pair; one spectrum did not distinguish the two sites, whereas the second favored T5 over S4 by Mascot site analysis (80.29% versus 16.17%). The site is therefore reported conservatively as S4/T5.

**Figure S7. *zhp-2(T210A)* mutants are competent for pairing and synapsis**
(A) Immunofluorescence micrographs of mid pachytene nuclei marked with SYP-1 (yellow) and HTP-3 (magenta). Scale bar, 5 μm. (B) Immunofluorescence micrographs of early pachytene nuclei marked with HIM-8 (yellow) and HTP-3 (magenta). Scale bar, 5 μm. (C) Immunofluorescence micrographs of *syp-1(T452A)* mutants marked with ZHP-2 (yellow) and HTP-3 (magenta) at the indicated stages. Scale bar, 5 μm. (D) Immunofluorescence micrographs of *syp-1(T452A)* mutants marked with PLK-2 (yellow) and HTP-3 (magenta) at the indicated stages. Scale bar, 5 μm.

**Figure S8. *zhp-2* mutants fail to form wild-type levels of double Holliday junctions**
(A) Super resolution micrographs of representative late pachytene nuclei marked with MSH-5 (yellow), COSA-1 (blue), and SYBR GOLD (magenta) for the indicated genotypes. Foci are enlarged and masks demonstrate singlets, doublets, and triplets. Scale bars, 1 μm. (B) Histogram showing the percentage of COSA-1 foci flanked by MSH-5 singlets, doublets, or triplets in the indicated genotypes. Number of foci scored (*n*), percentage of singlets, doublets, triplets: *ollas::cosa-1; gfp::msh-5* (216), 83.3, 15.3, 1.4; *ollas::cosa-1; gfp::msh-5; zhp-2(vv84)* (1007), 94.6, 5.0, 0.4; *ollas::cosa-1; gfp::msh-5; zhp-2(T210A)* (914), 95.3, 4.7.

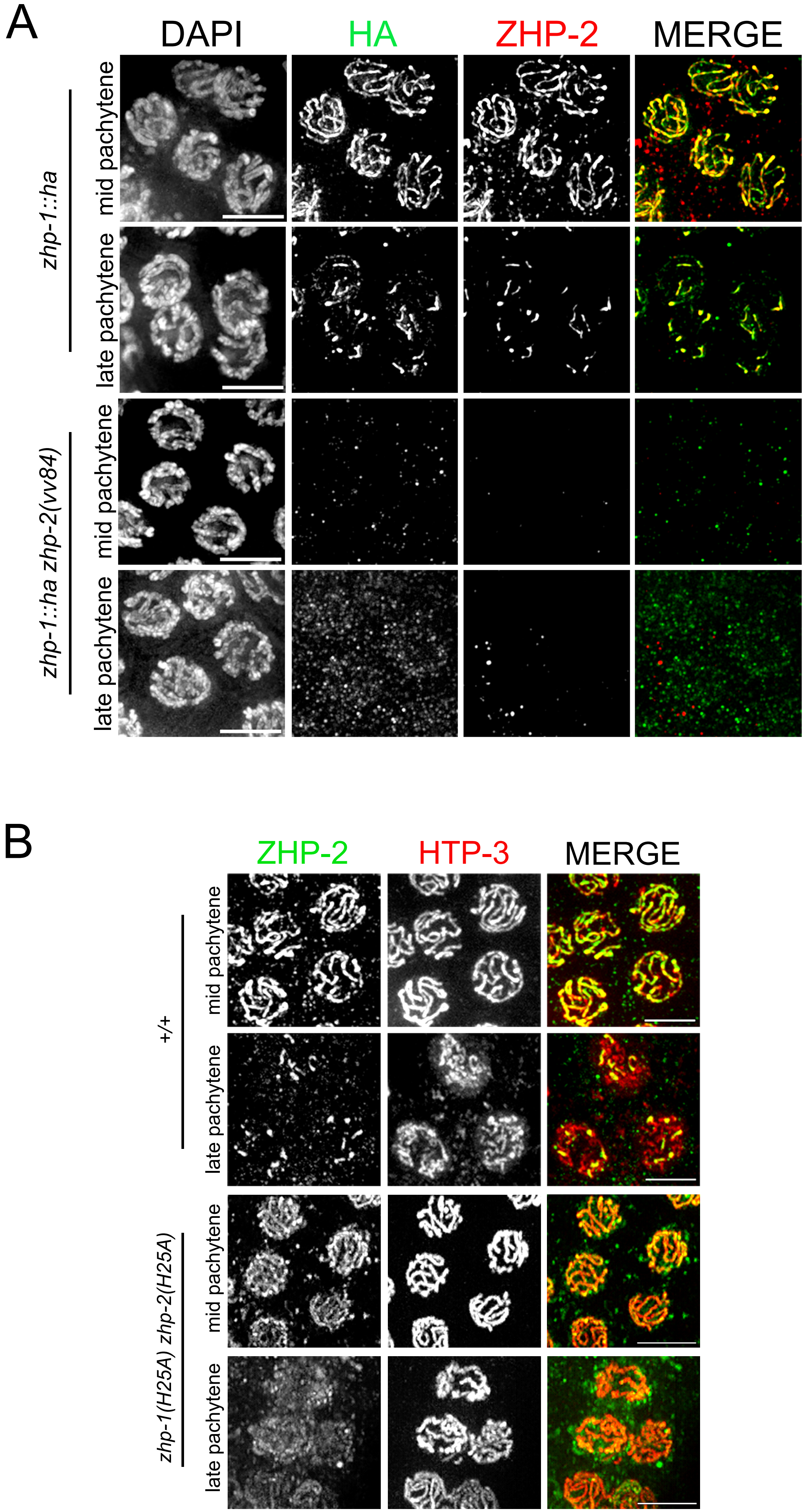

A
DAPI
HA
ZHP-2
MERGE
zhp-1::ha
mid pachytene
late pachytene
zhp-1::ha zhp-2(vv84)
mid pachytene
late pachytene
B
ZHP-2
HTP-3
MERGE
+/+
mid pachytene
late pachytene
zhp-1(H25A) zhp-2(H25A)
mid pachytene
late pachytene

Figure S2

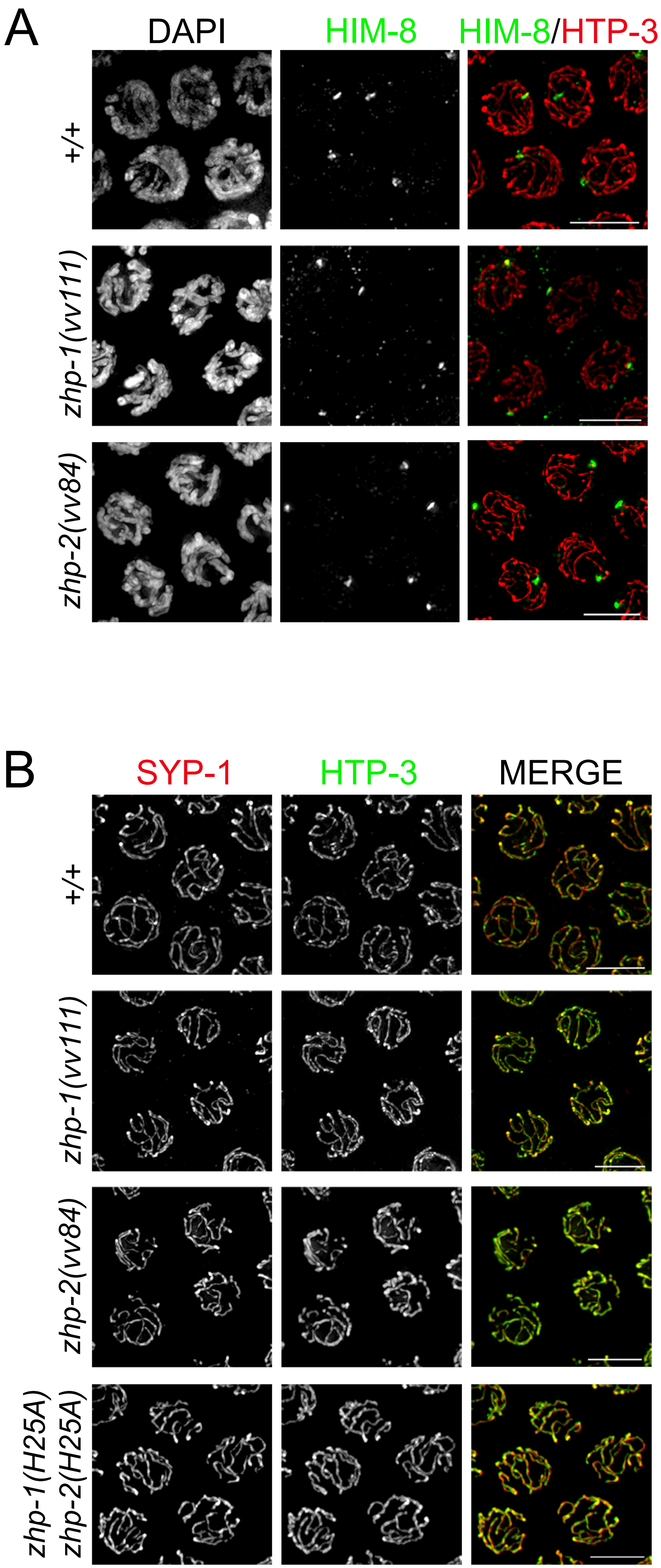
A
DAPI
HIM-8
HIM-8/HTP-3
+/+
zhp-1(vv111)
zhp-2(vv84)
B
SYP-1
HTP-3
MERGE
+/+
zhp-1(vv111)
zhp-2(vv84)
zhp-1(H25A)
zhp-2(H25A)

Figure S3

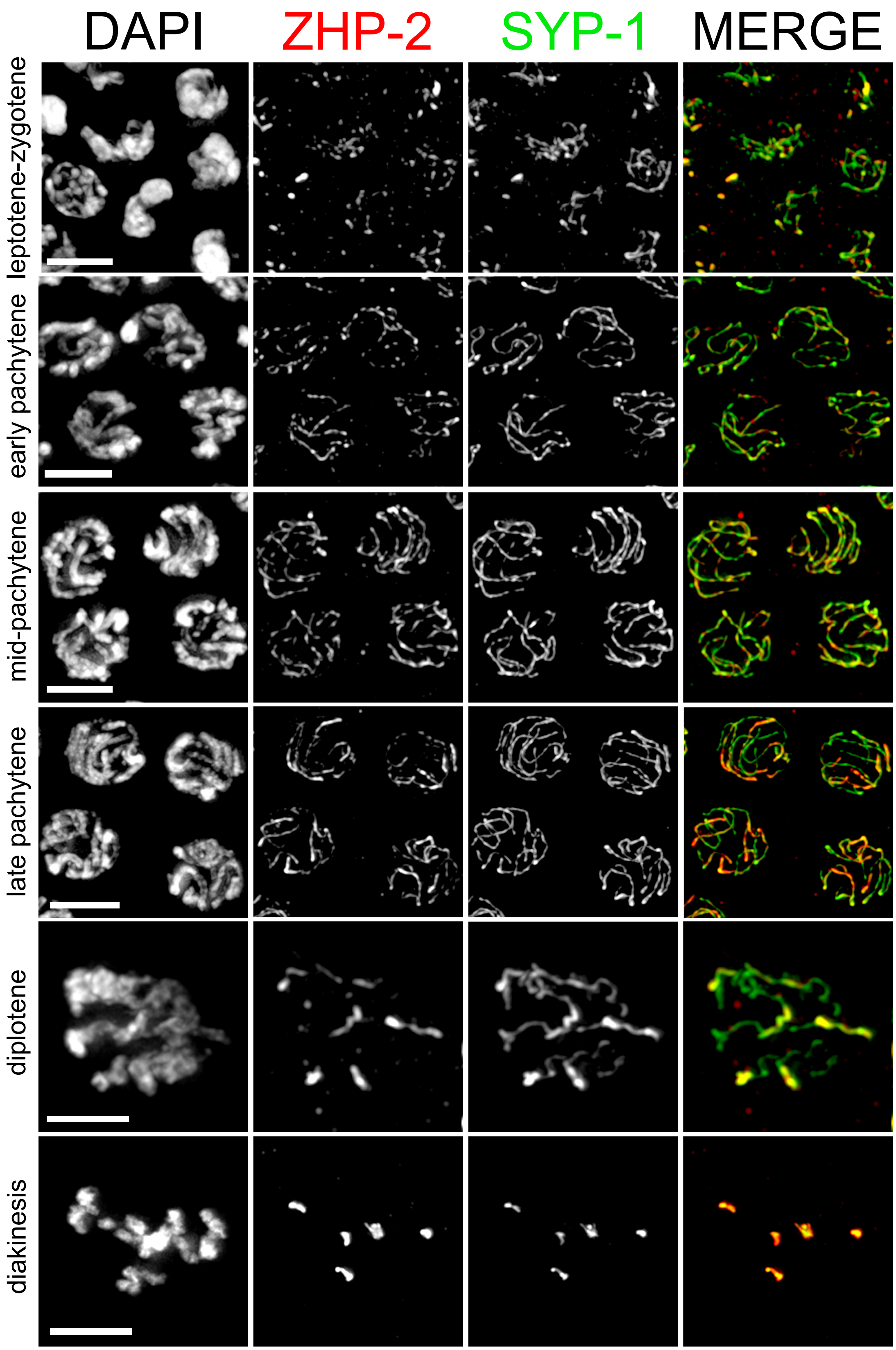
DAPI
ZHP-2
SYP-1
MERGE
leptotene-zygotene
early pachytene
mid-pachytene
late pachytene
diplotene
diakinesis

Fig. S4

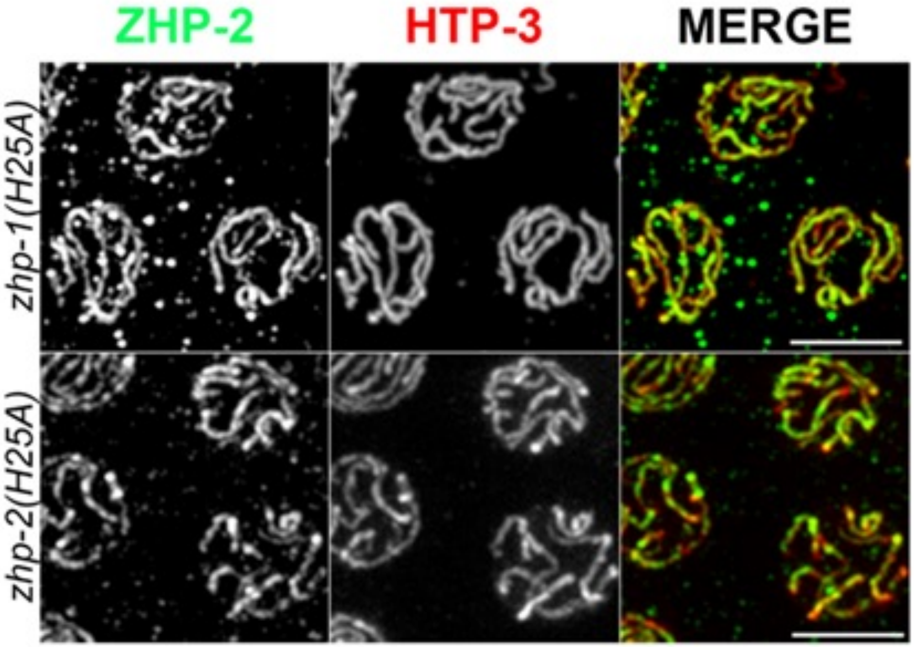

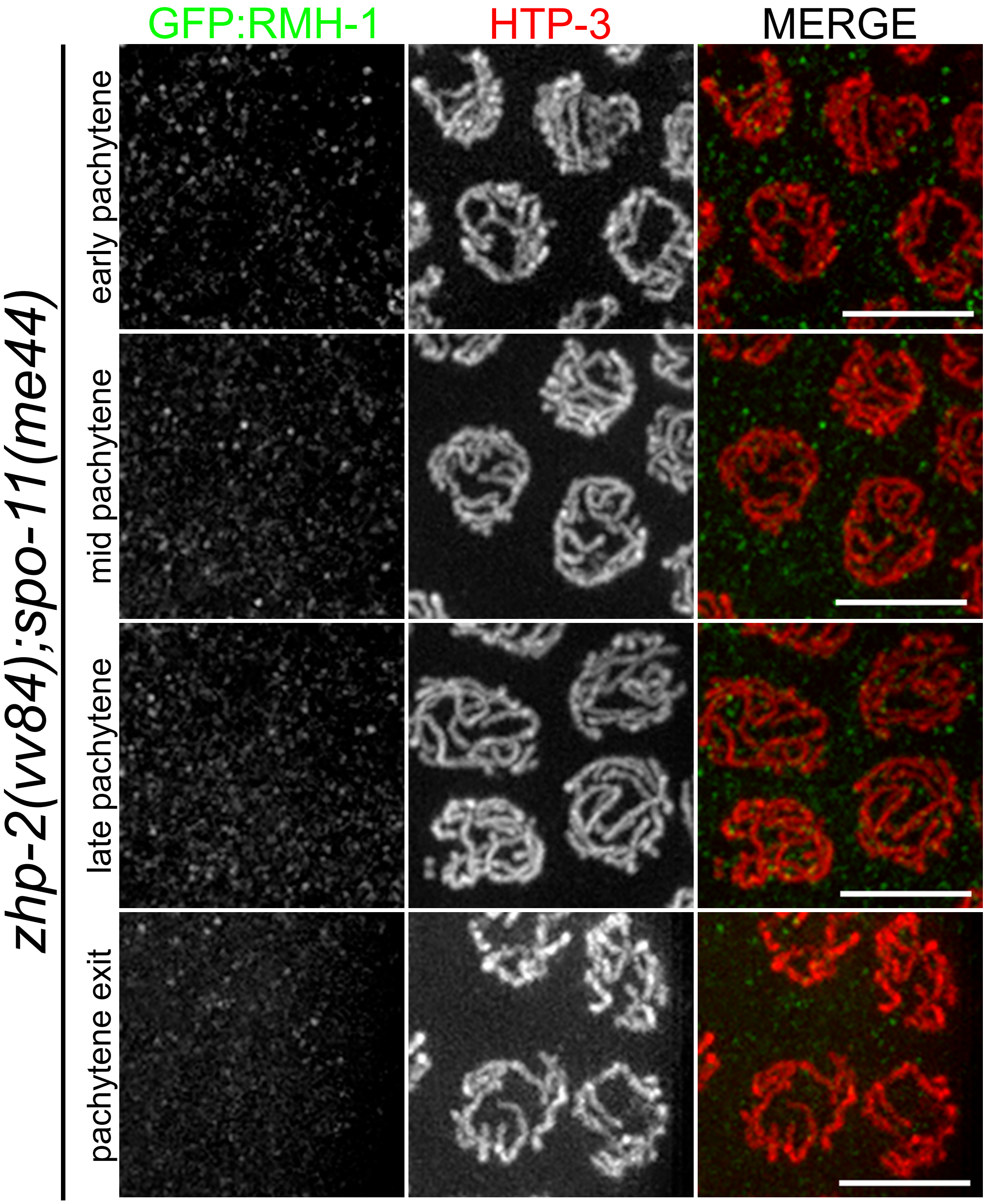
GFP:RMH-1
HTP-3
MERGE
zhp-2(vv84);spo-11(me44)
early pachytene
mid pachytene
late pachytene
pachytene exit

Figure S6

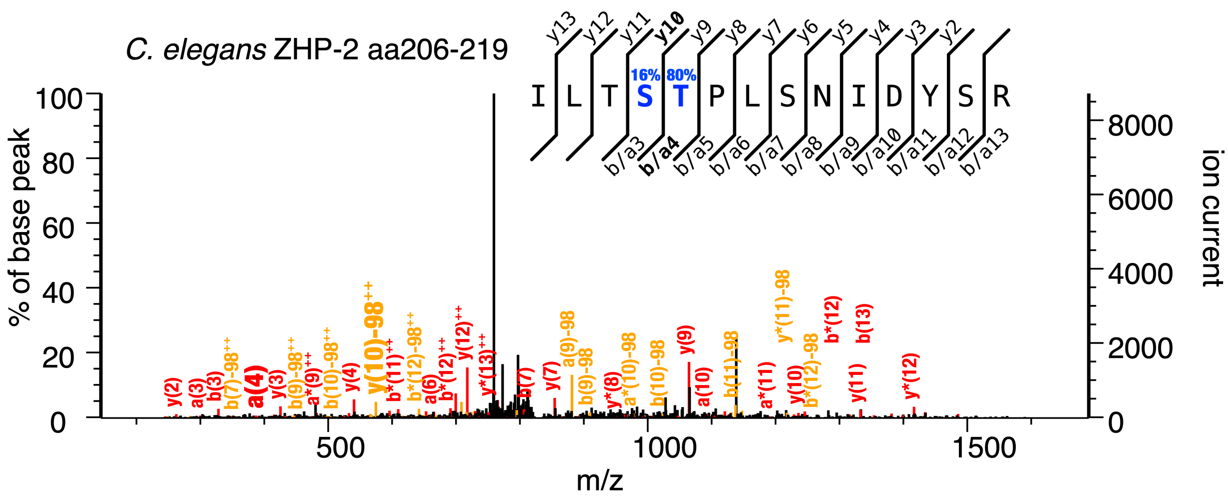

C. elegans ZHP-2 aa206-219
I L T S T P L S N I D Y S R
16% 80%
% of base peak
ion current
m/z

Figure S7

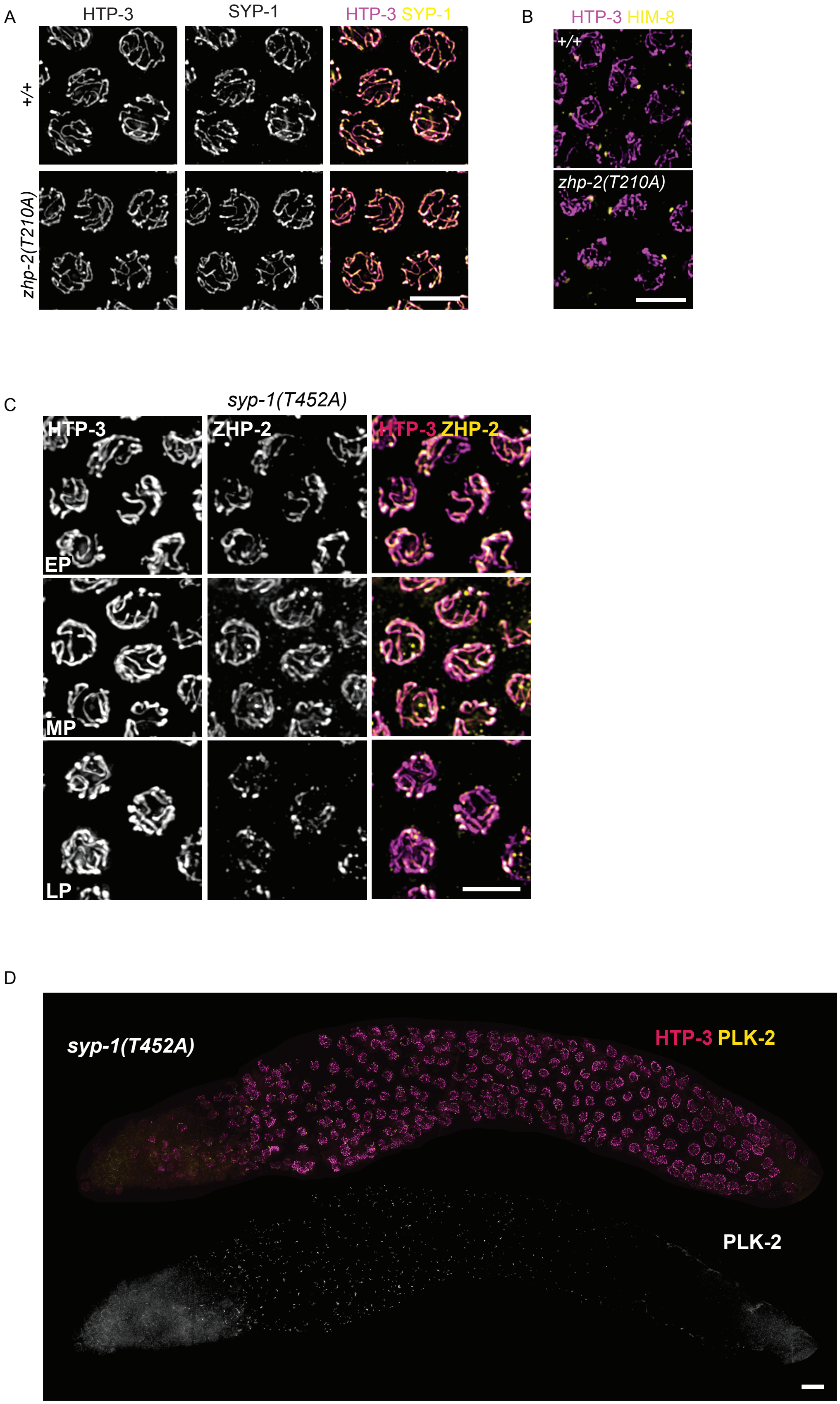
A
HTP-3
SYP-1
HTP-3 SYP-1
+/+
zhp-2(T210A)
B
HTP-3 HIM-8
+/+
zhp-2(T210A)
C
syp-1(T452A)
HTP-3
ZHP-2
HTP-3 ZHP-2
EP
MP
LP
D
syp-1(T452A)
HTP-3 PLK-2
PLK-2

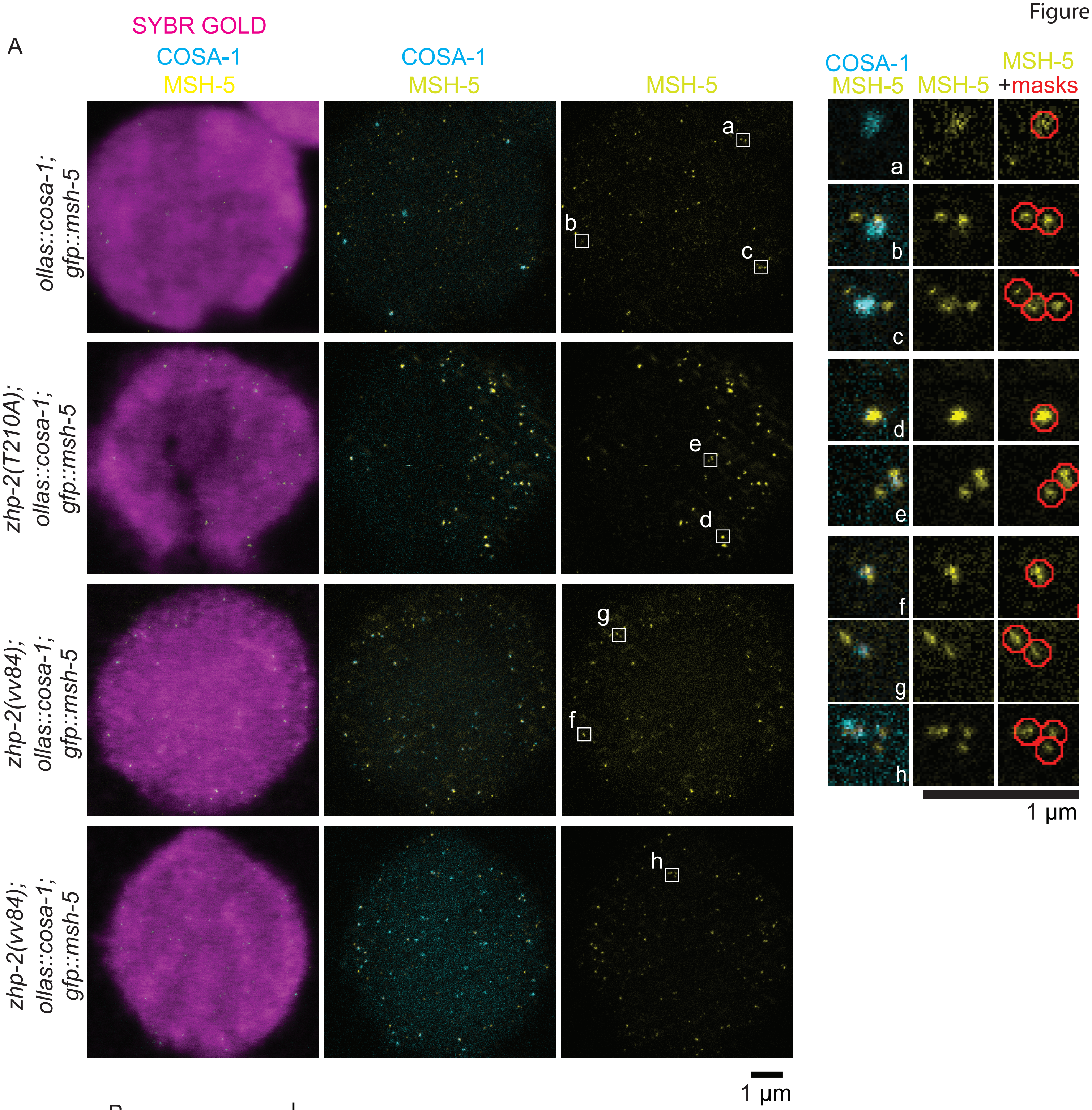
A
SYBR GOLD
COSA-1
MSH-5
COSA-1
MSH-5
MSH-5
COSA-1
MSH-5
MSH-5
MSH-5
+masks
ollas::cosa-1; gfp::msh-5
zhp-2(T210A); ollas::cosa-1; gfp::msh-5
zhp-2(vv84); ollas::cosa-1; gfp::msh-5
zhp-2(vv84); ollas::cosa-1; gfp::msh-5
a
b
c
d
e
f
g
h
1 µm
1 µm

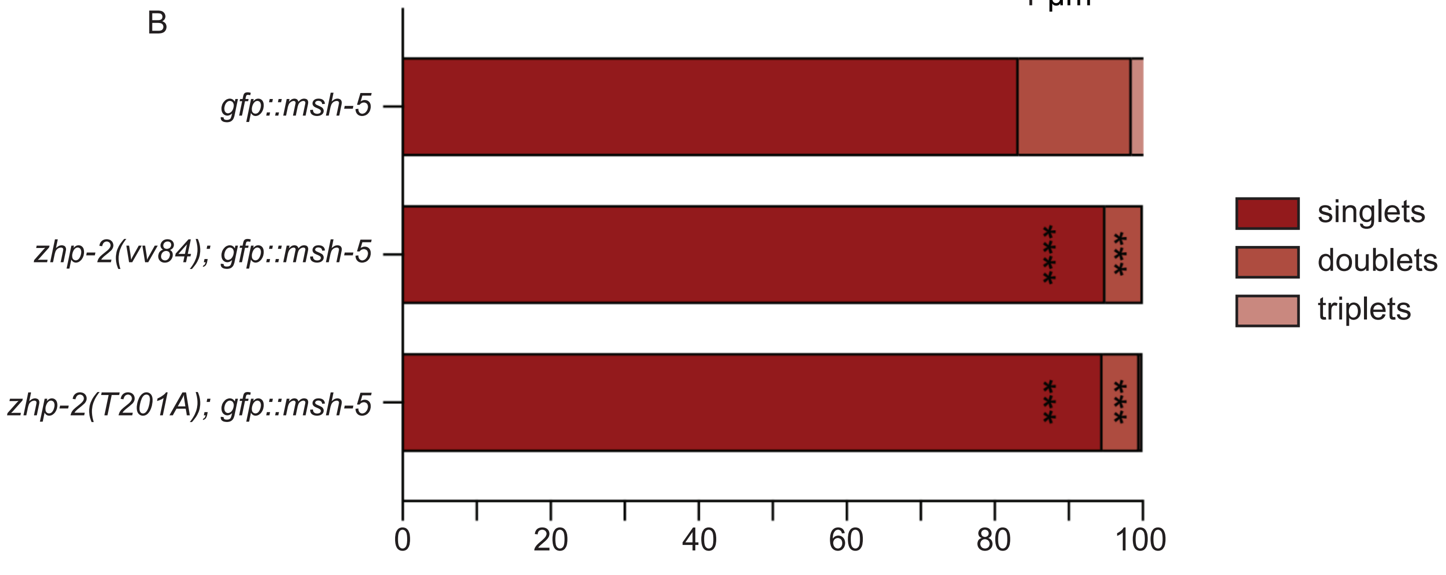
B
gfp::msh-5
zhp-2(vv84); gfp::msh-5
zhp-2(T201A); gfp::msh-5
****
***
***
***
0
20
40
60
80
100
singlets
doublets
triplets
